# 10.5 Tesla High-Resolution Macaque Brain MRI for In vivo and Ex vivo Connectivity Studies

**DOI:** 10.64898/2025.12.22.695917

**Authors:** Shaun Warrington, Mohamed Kotb Selim, Benjamin C. Tendler, Steen Moeller, Hamza Farooq, Wenchuan Wu, Pramod Pisharady, Gregor Adriany, Edward J. Auerbach, Davide Folloni, Alexander Bratch, Ana M.G. Manea, Taylor Grafft, Steve Jungst, Noam Harel, Matt Waks, Franco Pestilli, Essa Yacoub, Christophe Lenglet, Kamil Ugurbil, Sarah R. Heilbronner, Karla L. Miller, Saad Jbabdi, Jan Zimmermann, Stamatios N. Sotiropoulos

## Abstract

Mapping brain connectivity in primates remains a major challenge due to difficulties in resolving microscopic white matter architecture, while maintaining whole-brain coverage. Increasing imaging spatial resolution is key for disambiguating fibre configurations within smaller anatomical volumes. Here, we present novel developments that allow high-resolution diffusion MRI of the macaque brain using one of the world’s highest-field human MRI scanners operating at 10.5 Tesla, allowing both in vivo and ex vivo macaque brain imaging. Our approach achieves very high spatial resolution across both tissue states, (up to 580 *μm*)^3^ in vivo and (300 *μm*)^3^ ex vivo, with diffusion weighting up to b = 6000 *s*/*mm*^2^. We detail methodological advances in data acquisition, image reconstruction, processing and whole-brain tractography that overcome critical challenges associated with ultra-high-field imaging. This work establishes a new framework for high-resolution in vivo and ex vivo neuroimaging of the NHP brain at 10.5 T using a human bore scanner, paving the way for subsequent analyses of brain connectivity across species and tissue states at unprecedented detail. The dataset, along with all processing pipelines, containerised workflows, and reusable web services, is openly shared to support reproducibility and future integration with microscopy for studying white matter microstructure and connections at the mesoscale.

## Introduction

Despite well-characterised challenges^1^, diffusion magnetic resonance imaging (dMRI) is the only method available for non-invasively mapping white matter (WM) microstructure and connectivity in living primate brains, including humans. Diffusion MRI tractography approaches allow reconstruction of the WM connections that integrate and mediate information flow between remote brain regions, giving rise to cognition and behaviour^2^.

A clear advantage of dMRI over microscopy techniques^3,4^ is its ability to image the intact whole-brain at high-throughput. This is key for capturing long-range brain connections that may traverse the full length/width of the brain, and for imaging brains at population scales^5^. There is, however, a trade-off between spatial coverage and resolution. DMRI operates at the *macroscale* (imaging at *mm*), however the axonal fibres and WM bundles we wish to map are micro/mesoscopic (i.e. *µm*), respectively. Furthermore, dMRI is indirect; neuronal microstructure is *inferred* from measurements that reflect how the diffusion of water molecules is hindered by cellular structures within tissue. Unmet challenges in dMRI and tract reconstruction stem from ambiguities in resolving micro/mesoscopic WM fibre patterns from indirect macroscopic observations.

A two-pronged approach can be envisaged to address this challenge. Firstly, pushing the limits of dMRI acquisition and processing strategies^6–8^ can improve imaging resolution and contrast, and hence make the mapping of water diffusion processes to WM fibre patterns more specific^9–13^. Approaches can include, for instance, taking advantage of the greater intrinsic signal available when scanning at high and ultra-high field (UHF) strengths^14–16^; or taking advantage of increased diffusion weighting made possible with higher performance and higher strength gradient systems^17–21^. Secondly, integrating information across multiple scales (i.e. imaging resolutions) within modalities^11,22^ and multiple independent modalities (e.g. MRI and microscopy) can guide in resolving ambiguities^23,24^. Tracer studies and microscopy, for instance, along with MRI on the same tissue, provide independently observed and direct measurements of WM, against which MRI approaches can be constrained, validated, optimised and calibrated.

The Center for Mesoscale Connectomics (CMC, https://mesoscale-connectivity.org/) is taking such a multi-faceted approach for improving white matter tractography in the human and non-human primate (NHP) (macaque) brain. The CMC follows a multi-modal, multi-scale and axon-centric approach, developing technologies to image the same brains using i) dense and sparse axonal tracing^25,26^, ii) whole-brain polarization-sensitive optical coherence tomography (PS-OCT)^3,4^ and iii) whole-brain in vivo and ex vivo dMRI at UHF strength^14^ of both the human and NHP, using one of the strongest field (10.5 T) human MRI scanners available. As the same brains are imaged using each of the above modalities, the developed approaches will deliver unique data for better understanding WM connections and microstructure, improving fibre orientation modelling and tractography across resolutions and species.

In this paper, we present high resolution diffusion MR imaging of the NHP brain at UHF using the prototype 10.5 T scanner at the Center for Magnetic Resonance Research (CMRR), University of Minnesota. The large bore of the 10.5 T MRI scanner enables the same NHPs to be imaged in vivo, as well as ex vivo. We detail end-to-end methodological advances, from data acquisition to whole-brain tractography that overcome critical challenges associated with ultra-high-field imaging, and showcase data quality and processing efficacy across both tissue states. Over several years of developments, our approach achieves data of highest reported resolutions for this field strength and scanner type: up to (580 *μm*)^3^ in vivo and (300 *μm*)^3^ ex vivo, with diffusion weighting up to b = 6000 s/*mm*^2^ ex vivo. These are higher or comparable to the highest NHP resolutions achieved before using preclinical systems^23,27–29^ (hence ex vivo only), while addressing challenges of diffusion MRI using a large human-bore 10.5 T scanner. The proposed setup enables high-resolution UHF neuroimaging of NHP and paves the way for subsequent human^30^ and comparative studies of brain connectivity at unprecedented detail, both in vivo and ex vivo. The datasets, along with all processing pipelines, containerised workflows, and reusable web services, are openly shared (https://doi.org/10.25663/brainlife.pub.62) to support reproducibility and future integration with microscopy for studying white matter microstructure and connections at the mesoscale.

## Results

Diffusion MR imaging of both in vivo and ex vivo brains provides complementary benefits, but also pose unique challenges. Advantages of ex vivo imaging include longer acquisition times than for in vivo and minimal sample motion, no physiological artefacts and potential for higher signal to noise ratio (SNR)^31^, enabling much higher imaging resolutions. However, signal properties are affected by the tissue state due to biological (i.e. agonal/postmortem processes, removal of blood) and processing aspects (e.g. tissue fixation, sample temperature) involved in ex vivo imaging. Fixed tissue has reduced transverse relaxation (T_2_) and is typically reported to have diffusion properties (apparent diffusion coefficient, ADC) of the order of 2-3 times lower than living tissue^32–34^. In addition, imaging at ultra-high field (UHF) strengths introduces extra challenges. Even if the intrinsic signal to noise ratio (SNR) increases with field strength in supra-linear ways^35^, the shorter T_2_ and increased susceptibility-induced distortions at UHF compared to low field increase the need for faster acquisition.

We considered these competing aspects for developing dMRI approaches of the NHP brain using a large bore human UHF (10.5 T) MRI scanner with a modest 70 mT/m gradient strength. This is a significantly more challenging task in terms of main magnetic field (B_0_), radiofrequency (RF) coil field (B_1_), and gradient uniformities compared to dMRI using high-gradient pre-clinical MRI systems. However, the use of a large-bore scanner enables the scanning of the same NHPs in vivo and ex vivo on the same system and carries the potential of translating developments more directly to UHF dMRI of the human brain. We addressed these challenges and developed optimised protocols for dMRI acquisition and end-to-end processing in the macaque brain, across both tissue states (Fig. 1). The in vivo acquisition relies on the use of accelerations, including echo planar imaging (EPI) and k-space undersampling, to mitigate for distortions and artefacts at UHF^14,36^. The ex vivo acquisition employs diffusion-weighted steady state free precession (DW-SSFP) imaging, which addresses the short T_2_ and low diffusivities of fixed tissues, whilst simultaneously enabling inherently low-distortion data, even at UHF, due to compatibility with a heavily segmented readout^31,37^. The development of custom-made UHF receive/transmit coils^38^ was of particular importance for the efficacy and success of these protocols, providing gains in SNR and signal homogeneity.

**Figure 1.**
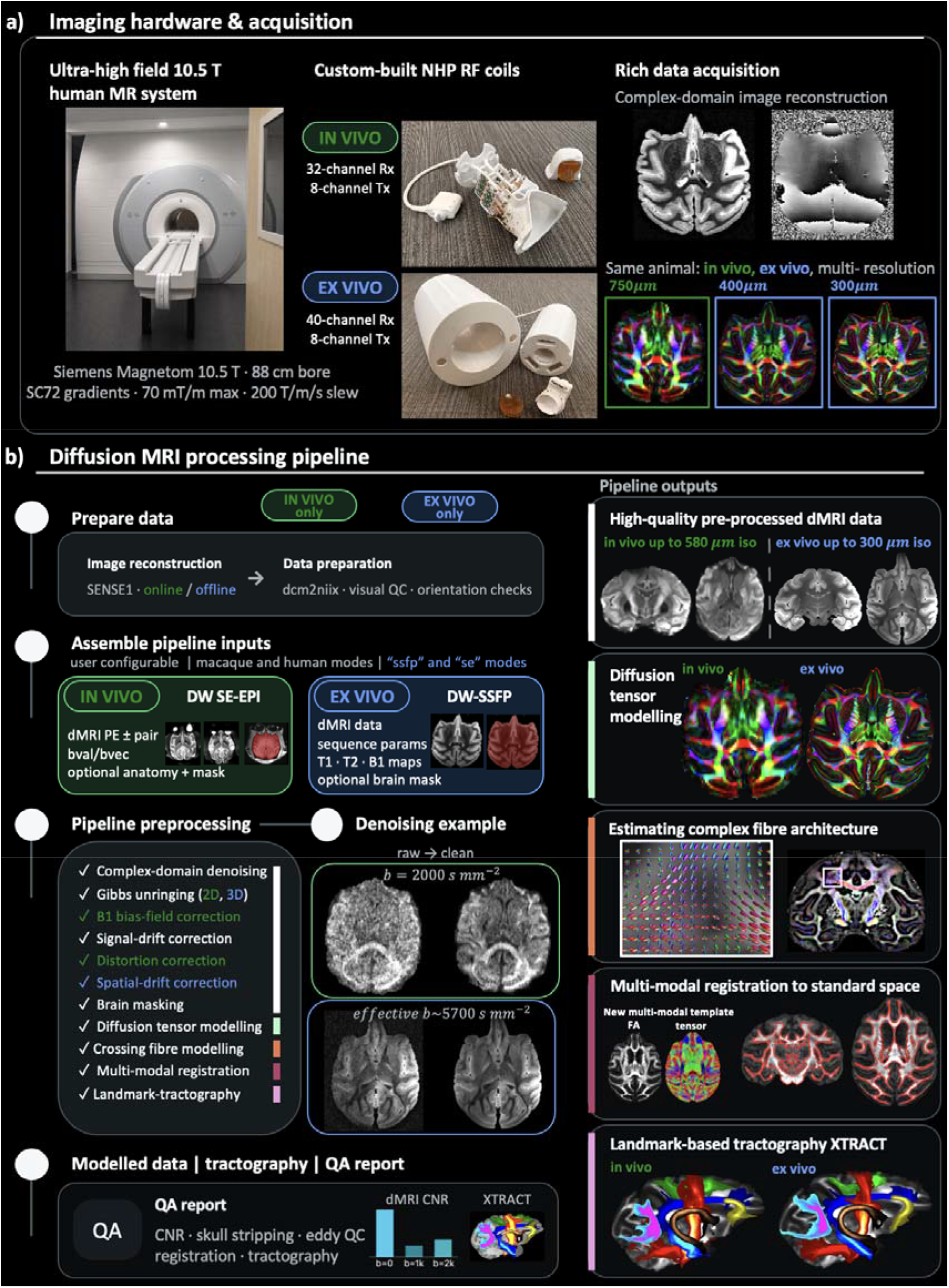
Overview of end-to-end acquisition and processing for in vivo and ex vivo diffusion MR imaging of the NHP at 10.5 T. a) dMRI data are acquired using the ultra-high field human scanner Siemens Magnetom at the CMRR using custom-made RF coils. Complex domain (magnitude and phase) data are acquired and the same animals are imaged both in vivo and ex vivo. b) The end-to-end processing pipelines for in vivo and ex vivo data, including denoising, distortion corrections where applicable, quality control, multi-modal registration, fibre orientation mapping and tractography (expanded version displayed in Supplementary Fig. S3). After image reconstruction and conversion to NifTI, the in vivo pipeline requires the dMRI data, a phase encoding (PE) blip-pair, text files containing the b-values (bvals) and diffusion gradient vectors (bvecs), and optional anatomical scan and brain mask for skull stripping. The ex vivo pipeline requires a single dMRI dataset (DW-SSFP) along with the coregistered DW-SSFP parameter maps (T1 map, B1 flip angle map, T2 map – see Methods), parameter files describing the repetition time (TR), flip angles, diffusion gradient amplitudes and durations and diffusion gradient vectors. Both pipelines are generalised to the human brain and the ex vivo pipeline also generalises to conventional spin-echo (DW SE-EPI) acquisitions. The pipelines produce quality assurance reports at several stages, which are summarised into a single PDF at the end of processing.

The different aspects of the acquisition and processing are presented in the following sections, showcasing data and tractography quality. The developments proposed here demonstrate, for the first time, feasibility for high-resolution in vivo and ex vivo dMRI of the same NHP brain at a human bore 10.5 T scanner, and pave the way for comparable UHF studies of human brain connectivity across both tissue states.

### End-to-end acquisition & processing for in vivo and ex vivo macaque dMRI tractography at 10.5 T

Using the prototype Siemens Magnetom 10.5 T human MRI scanner at the CMRR, University of Minnesota (Fig. 1a), we have imaged to date six NHPs (rhesus and cynomolgus macaques), five of those in vivo, and two of those ex vivo (Supplementary Table S1). Acquisition protocols and hardware (RF coils and gradients) were optimised through an iterative process (Supplementary Figs. S1, S2), balancing spatial and angular contrast/resolution, geometric distortions, signal homogeneity, contrast to noise ratio (CNR), acquisition time, and whole-brain coverage whilst managing the engineering challenges associated with using a prototype large bore (88 cm) UHF MR system. Full details on ex vivo acquisition optimisation can be found in^31^.

Anesthetised macaques were imaged using a pulsed gradient spin echo, diffusion-weighted echo planar imaging (EPI) sequence (DW SE-EPI) with two shells (b=1000, 2000 s/mm^2^), 54 volumes per shell, four repeats and pairs of acquisitions with reversed phase-encoding directions were acquired over approximately 2.5 hours (Table 1, Supplementary Table S2). Ex vivo, a single-line readout DW-SSFP sequence with two shells (equivalent b-value of approximately 3,900 and 5,900 s/mm^2^) across 121 directions was used, acquired over approximately 54 hours (Table 1, Supplementary Table S3). We were able to achieve isotropic spatial resolutions of up to (580 *μm*)^3^ in vivo and (300 *μm*)^3^ ex vivo, with the first NHPs imaged at (750 *μm*)^3^ in vivo and (400 *μm*)^3^ ex vivo. For both in vivo and ex vivo, complex data (magnitude and phase) were saved, enabling complex-domain denoising and offline reconstruction.

**Table 1.** Overview of key acquisition parameters by imaging resolution and state. TE = echo time; TR = repetition time; ES = echo spacing

| | Spatial res<br>($\mu\text{m}$ iso) | TE<br>(ms) | TR<br>(s) | ES<br>(ms) | b-values<br>($\text{s}/\text{mm}^2$) | #volumes<br>per shell | #repeats | Scan<br>time |
| --- | --- | --- | --- | --- | --- | --- | --- | --- |
| 2D DW<br>SE-EPI | 750 | 66 | ~8.5 | 0.33 | 0, 1000,<br>2000 | 8, 54, 53<br>(920 total) | 8<br>(4 AP, 4 PA) | ~2 hrs |
| 2D DW<br>SE-EPI | 580 | 79 | ~9.2 | 0.40 | 0, 1000,<br>2000 | 8, 54, 53<br>(920 total) | 8<br>(4 AP, 4 PA) | ~2.5<br>hrs |
| 3D DW-SSFP | 400 | 21 | 0.026 | - | 0, 2900,<br>5700 | 8, 60, 60<br>(128 total) | - | ~28<br>hrs |
| 3D DW-SSFP | 300 | 21 | 0.027 | - | 0, 3900,<br>5900 | 8, 60, 60<br>(128 total) | - | ~54<br>hrs |

We developed end-to-end processing (Fig. 1b, Supplementary Fig. S3), from image reconstruction to whole brain tractography, optimised to brain state (in/ex vivo) and imaging at 10.5 T. Indicative pipeline step developments are shown, including: i) A custom CMC image reconstruction, using a SENSE1 coil-combination^39^ to increase the signal dynamic range (Supplementary Fig. S4), followed by complex domain denoising^40^ (Supplementary Fig. S5), to further reduce noise-induced variance and biases^41^. ii) Optimised preprocessing including Gibbs unringing^42–44^ (Supplementary Fig. S6), signal drift correction (Supplementary Fig. S7), high-order (cubic) and advanced distortion correction^45,46^ for susceptibility and eddy-current induced distortions and motion and their interactions when imaging with accelerated acquisition strategies in vivo, and quality control. iii) Newly developed multi-modal NHP templates and approaches for accurate multi-modal (anatomical and dMRI) registration^47^ to the NIMH Macaque Template (NMTv2) space. iv) Fibre orientation reconstruction approaches using both DW SE-EPI (in vivo) and DW-SSFP (ex vivo) data and up to three orientations per voxel, extending previous work^48,49^. v) Cross-species (macaque and human) landmark-based tractography with protocols in NMT space^50,51^ to enable future comparative studies. vi) Newly developed tractography protocols for reconstructing the stria terminalis and fornix, optimised for and with high resolution ex vivo imaging, extended to in vivo imaging. These showcased the benefit of high resolution for tractography in localising relatively small nearby tracts, extending also to examples delineating the external/extreme capsules, and the amygdalofugal tract running nearby the anterior commissure and the uncinate fasciculus.

Workflows are available for bare metal installation or through containerised environments and employ a hashing and quality assurance (QA) system to ensure efficient and robust data processing. In addition, workflows are integrated into brainlife.io^52^ to enable decentralised cloud-based processing. Through minimal tweaking of a configuration file, the pipelines are generalisable to NHP and human data, and the ex vivo pipeline can operate on either conventional DW-SE single-line readout acquisitions or DW-SSFP single-line readout acquisitions, as done for the current data. The current data are also publicly available on brainlife.io (https://doi.org/10.25663/brainlife.pub.62) and will be further augmented by UHF data from additional macaques, as well as from humans, including both in vivo and ex vivo MRI, PS-OCT and tracing data.

### High-order and UHF-optimised distortion correction for optimising in vivo preprocessing efficacy

To mitigate the greater susceptibility-induced distortions at UHF, which can be severe for in vivo DW SE-EPI data, we identified optimal solutions that firstly included a high in-plane acceleration factor (GRAPPA=3), along with iterative B_0_ shimming and power calibrations for each animal using a transmit B_1_ fieldmap. Secondly, we used blip-reversed *b=0* acquisitions^45^ to estimate an off-resonance fieldmap and correct for geometric distortions (Fig. 2a). The efficacy of distortion correction was assessed for two different phase encoding directions (AP/PA vs LR/RL), with the former showing better alignment with a distortion-free anatomical (red boundaries, Fig. 2b). This was likely caused by the larger proportion of fat and muscle relative to brain/head size, which is distorted along LR/RL compared to AP/PA direction. As fat and muscle proportion is highly variable across species of macaque and sexes, and is typically much greater than in the human, we chose the AP/PA phase-encoding direction to provide more generalisable performance across animals.

**Figure 2.**
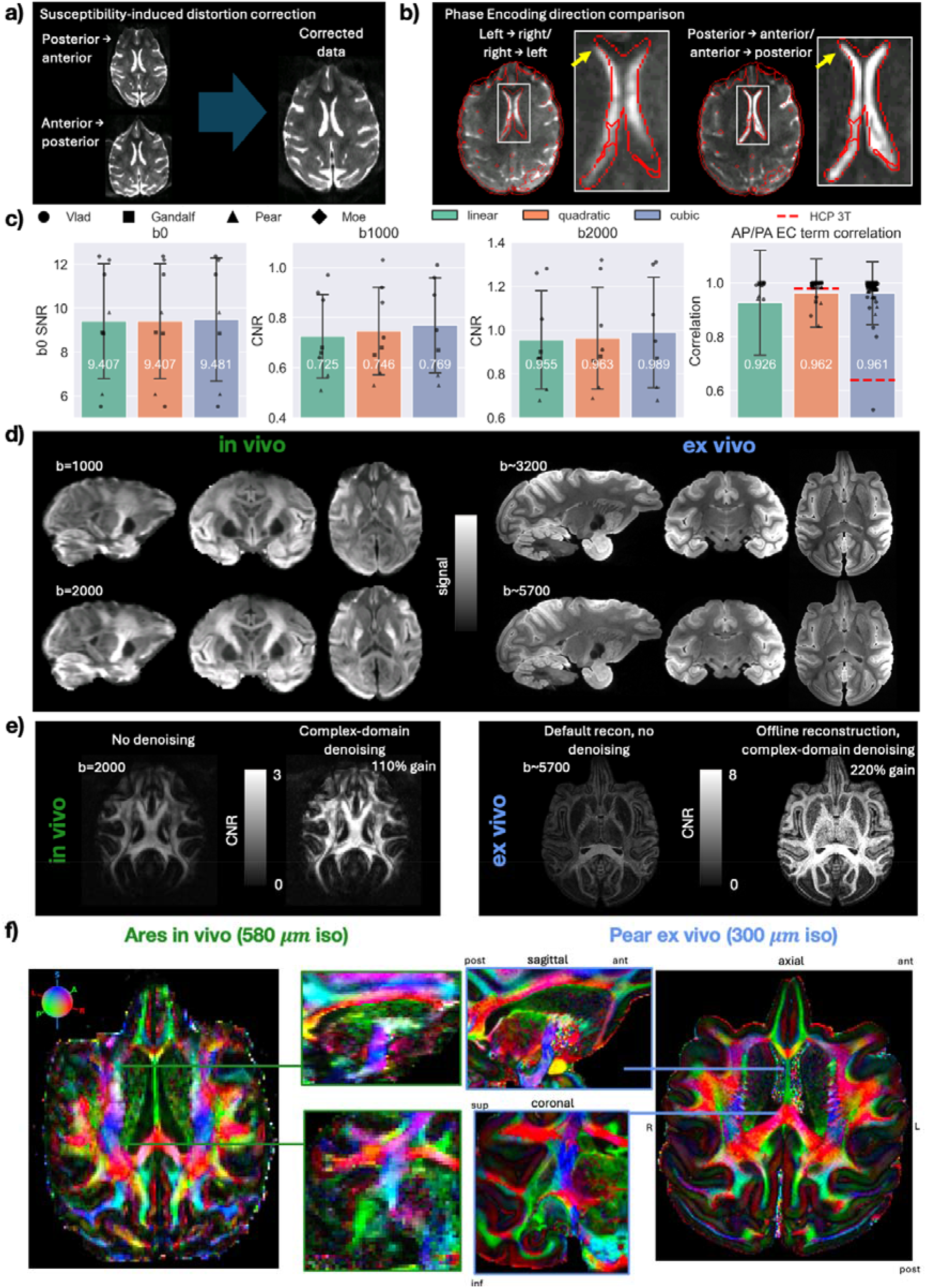
Optimised distortion correction for increasing preprocessing efficacy of ultra-high field dMRI. a) Correction of geometric distortions induced by susceptibility off-resonance. For in vivo data, pairs of reversed phase encoding data are acquired to estimate B_0_ off-resonance fieldmaps in dMRI space. b) Comparisons of susceptibility-distortion correction efficacy using left-right vs anterior-posterior phase encoding. Post-distortion correction, dMRI data are registered to T2w anatomical (i.e. distortion-free) space and the red line outlines the tissue boundaries derived from the T2w image. c) Quantitative assessment of eddy current models, comparing linear (green), quadratic (orange) and cubic (blue) eddy current models. Metrics include tSNR (b=0 shell only, left-hand plot), angular CNR across b-shells (middle plots) and the correlation (right-hand plot) between estimated eddy current model parameters across independent AP-PA runs. Values are derived from the four in vivo acquisitions. Red dashed lines indicate correlation^46^ “thresholds” for quadratic and cubic models applied to lower field HCP 3T data. d) Average images per b-value for in vivo and ex vivo data, following distortion correction. e) Examples of CNR gains for in vivo at 750 µm (left) and ex vivo at 300 µm (right) when comparing default scanner reconstruction with no denoising and custom offline reconstruction with complex-domain denoising (both following distortion correction). f) Whole-brain colour coded fractional anisotropy maps for Ares 580 µm in vivo (left) and Pear 300 µm ex vivo (right), highlighting preprocessing efficacy for both tissue states and detailing gains through high resolution ex vivo imaging.

We subsequently explored optimal eddy current models for data acquired at this field strength. At lower field, linear or quadratic spatial models suffice for capturing the off-resonance field generated by eddy currents^46^. In fact, higher-order cubic models overfit to data and cannot be reliably estimated. For these 10.5 T datasets, we found that cubic model estimation was supported and it improved distortion correction (Fig. 2c, Supplementary Figs. S8, S9). Post-correction, b=0 SNR was approximately 0.7% higher and angular CNR values were 3-6% higher with a cubic model compared to lower order models, suggesting less residual distortion and better distortion correction efficacy (Fig. 2c, left). At the same time, cubic model parameters estimated from two independent runs on the same brain (AP vs PA) were almost perfectly correlated (Fig. 2c, right), suggesting that the cubic model captured genuine eddy current off-resonance effects (for comparison, correlation of quadratic and cubic model parameters on HCP 3T data are shown with the dashed red lines, indicating less support for higher order models at lower field). Finally, we used automatic outlier detection and replacement^46^, estimated and corrected the susceptibility with motion interactions, yielding optimal eddy correction for ultra-high field distortions (Supplementary Figs. S8, S9).

Following image reconstruction, denoising and distortion correction, angular CNR was improved approximately 110% (i.e. doubling) for in vivo and 220% (i.e. tripling) for ex vivo using our optimised pipeline, while maintaining signal at a whole-brain level (Fig. 2d,e), speaking to the preprocessing efficacy. This can be further appreciated in the quality of DTI-derived maps (Supplementary Fig. S10) and RGB colour-coded fractional anisotropy maps (Fig. 2f). These demonstrate preprocessing efficacy for example data imaged in both tissue states, and the enhanced detail gained when imaging using the higher resolution DW-SSFP technique.

### Multi-modal registration enabled through the development of a new NHP template

Given the CMC aims for MRI and microscopy integration, alignment of different datasets is of paramount importance. To enable optimal alignment across tissue states, and with the NMT macaque template space^53^, we developed a new multi-modal (anatomical and dMRI) NMT-space template and used the recently developed multi-modal non-linear registration framework (MMORF)^47^. This enabled us to take advantage of complementary information from dMRI diffusion tensors and scalar maps (fractional anisotropy, FA) to drive the estimation of native to template space transformation fields more accurately. NMT-space diffusion templates (Fig. 3a) were derived through an iterative multi-subject alignment and refinement procedure using an external dataset of six ex vivo macaque brains.

**Figure 3.**
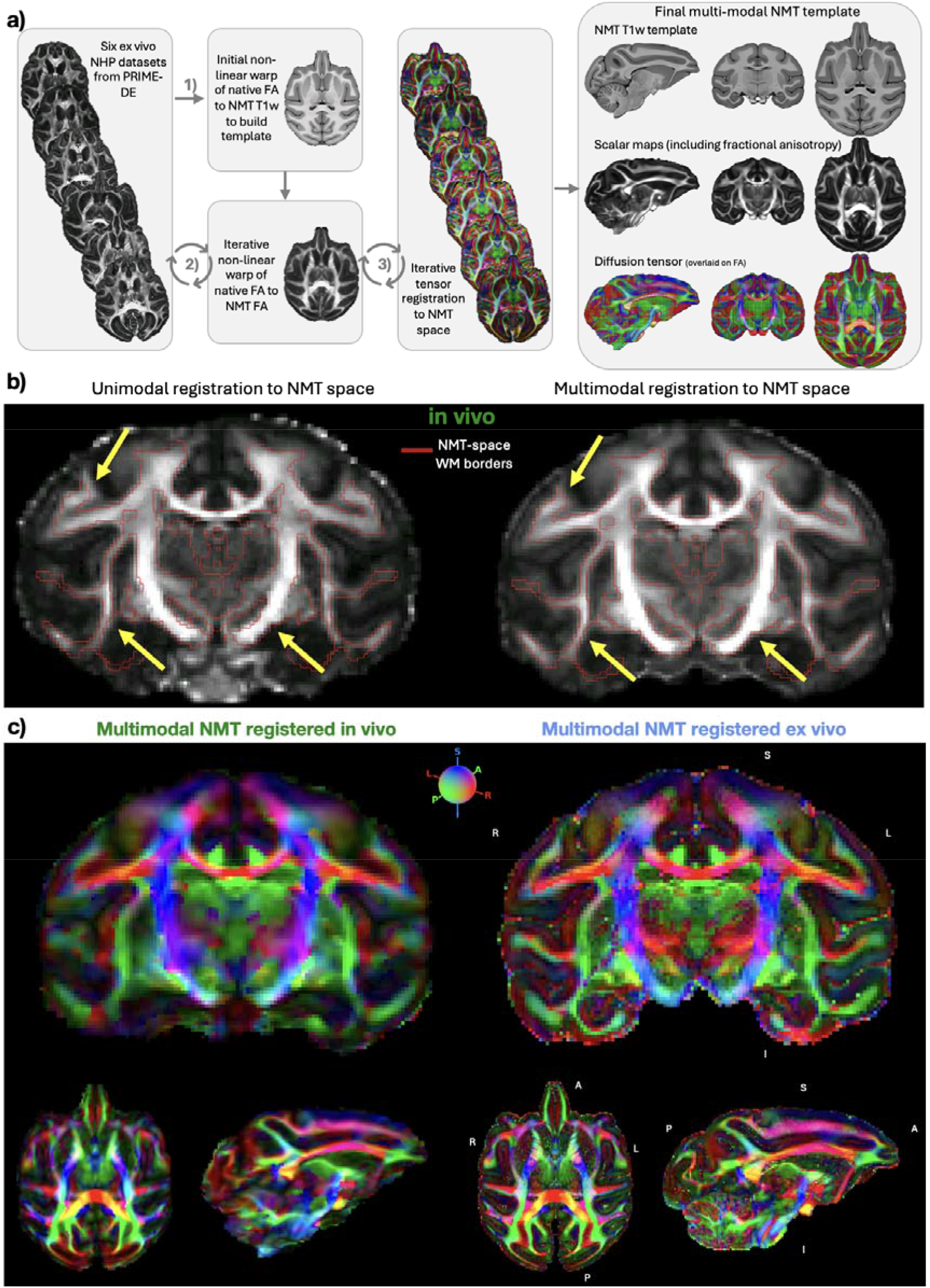
Development of a new multi-modal NMT-space template facilitating improved cross-state anatomical alignments and comparisons. a) Multimodal (anatomical and diffusion tensors) template building. Diffusion tensor metrics (fractional anisotropy, FA) were registered to NMT space through a conventional non-linear approach and a cohort-specific template was derived through iterative refinement. Native space tensors were finally transformed into NMT space with appropriate tensor rotations before being decomposed into diffusion tensor maps, including FA and mean diffusivity (MD) and fibre orientation maps. b) Comparison of FA maps of the same brain aligned in NMT space through unimodal (i.e. conventional) and multimodal registrations. Red outlines indicate the NMT template WM/FA edges (FA threshold of 0.2). Yellow arrows indicate examples of misalignment that the unimodal registration suffers from, which are resolved with the multimodal registration approach. c) In vivo and ex vivo FA maps of the same brain aligned in NMT space and colour-coded by the principal fibre orientation estimated from diffusion tensor modelling. Note: whilst registration enables direct comparisons in the same image space, some spatial blurring is induced through interpolation.

Using the constructed FA and diffusion tensor macaque template, in vivo and ex vivo data could be aligned to the template space using multimodal non-linear registration via their FA and tensor images. The multimodal registration achieved superior alignment to the NMT space compared to conventional unimodal (FA-based) non-linear registrations (Fig. 3b). This was true for both in vivo and ex vivo data, allowing direct and accurate comparison of the same animal imaged across tissue states (Supplementary Fig. S11) and facilitating downstream registration-reliant landmark-based tractography. NMT-registered RGB-orientation-coded FA maps demonstrate the efficacy in the alignment of both tissue states, as well as the benefit of higher resolution for the ex vivo DW-SSFP data (Fig. 3c), even in the event of spatial blurring due to interpolations during registration. Notably, the high agreement between template-space, in vivo and ex vivo data is also indicative that the CMC pre-processing steps successfully resolve the majority of geometric distortions in vivo.

### DW SE-EPI & DW-SSFP fibre orientation mapping enables WM tractography in the in vivo and ex vivo brain

Even if plenty of approaches exist for spherical deconvolution of (multi-shell) diffusion-weighted spin echo (DW SE) dMRI data, fibre orientation mapping is less straightforward for DW-SSFP data, where the signal has T1, T2, B1 dependences and diffusion-weighting does not correspond to a single well-defined b-value^54^ (i.e. definition of “b-shells” can only be approximate). Building upon previous DW-SSFP models^48,49^, we extended modelling approaches to reconstruct fibre crossings and fibre orientation distributions (FODs) using parametric deconvolution^55^ both in vivo (DW SE-EPI, Fig. 4a left) and ex vivo (DW-SSFP) (Fig. 4a middle). Multi-way crossings could be consistently mapped (up to three orientations shown in Fig. 4a), providing evidence for the data angular contrast and preprocessing efficacy across both tissue states. Notably, using deconvolution of DW-SSFP data against the conventional DW-SE fibre response kernel and with an effective b-value approximation^56^ provided suboptimal results compared to using an SSFP dedicated kernel and model that does not require such approximations (Supplementary Fig. S12). This highlights the value of the developments and feasibility of using DW-SSFP data for multicompartment microstructure modelling. We further confirmed that FODs could be obtained from the ex vivo DW-SSFP data using non-parametric multi-shell multi-tissue (MSMT) constrained spherical deconvolution (CSD) (Fig. 4a right), when similarly relying on an effective b-value approximation^56^.

**Figure 4.**
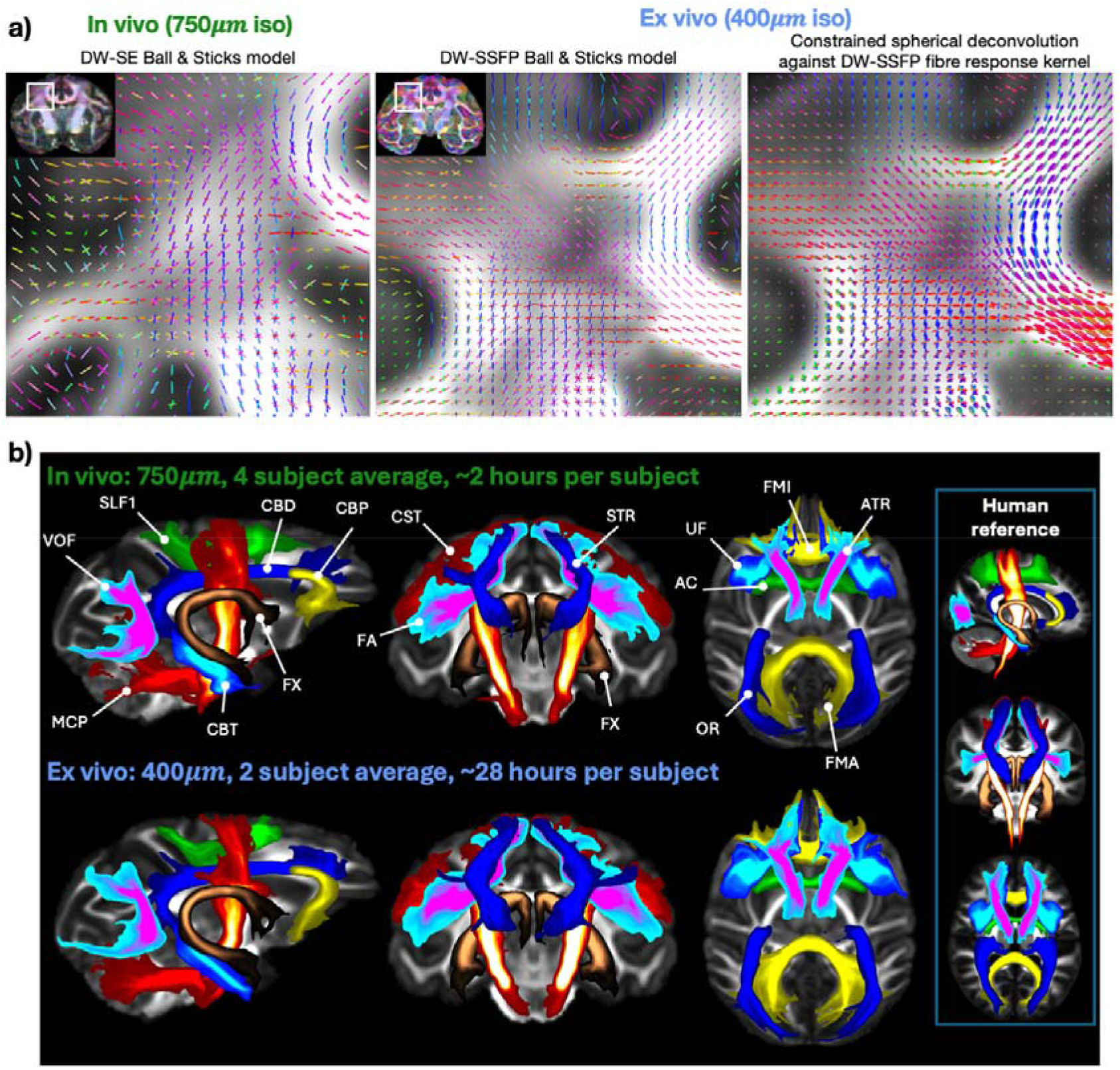
Fibre orientation estimation and landmark-based tractography for the in-vivo and ex-vivo NHP brain at 10.5 T. a) Fibre orientation estimations for in vivo data (left), using the conventional spin-echo (DW-SE) diffusion kernel with the ball and three-sticks model, and ex vivo data, using a DW-SSFP kernel with a modified ball and three-sticks model (middle) and using multi-shell multi-tissue (MSMT) constrained spherical deconvolution against a single fibre response learnt from the data (right). b) Examples of reconstructions of major white matter fibre bundles through landmark-based probabilistic tractography. Group-averaged maximum intensity projections for in vivo (top, averaged across four animals) and ex vivo (bottom, averaged across two animals) acquisitions and reference tract atlases for the human brain from^51^, averaged across 1,065 Human Connectome Project (HCP) participants. AC: anterior commissure, ATR: anterior thalamic radiation, CBD: dorsal bundle of the cingulum, CBP: peri-genual bundle of the cingulum, CBT: temporal bundle of the cingulum, CST: cortico-spinal tract, FA: frontal aslant tract, FMA: forceps major, FMI: forceps minor, FX: fornix, MCP: middle cerebellar peduncle, OR: optic radiation, SLF1: the first branch of the superior longitudinal fasciculus, STR: superior thalamic radiation, UF: uncinate fasciculus, VOF: vertical occipital fasciculus.

Tractography based on NMT-space anatomical landmarks^50,51^ allowed reconstruction of major white matter fibre bundles from each dataset, including projection, commissural, association and limbic bundles (full list of tracts in Supplementary Table S4) across the whole macaque brain (Fig. 4b). Examples of a subset of these reconstructions for both in vivo and ex vivo data averaged across animals (4 animals in vivo at (750 *μm*)^3^, 2 animals ex vivo at (400 *μm*)^3^) reveal correspondence in bundle reconstructions across states/resolutions. Median correlation between in vivo and ex vivo averages was 0.63 for all tracts and 0.66, 0.64, 0.64, 0.62, 0.61 for association, projection, commissural, limbic and subcortical bundles respectively; showing slightly higher agreement in association and lower agreement in subcortical bundles across resolutions. Notably, all the tractography protocols have been designed in our previous work to be generalisable to humans^51,57^ (exampled in Fig. 4b), enabling subsequent cross-species comparisons and integration^58^.

### High spatial resolution improves tractography of nearby subcortical white matter bundles

Having established that both our in vivo and ex vivo data can yield tractography reconstructions of major white matter bundles across the whole brain, we explored how spatial resolution differences can benefit tractography of small white-matter bundles that pass through compact regions of dense fibre organisation^12^. Indeed, a primary advantage of ex vivo diffusion MRI is the ability to use long acquisition times with minimal motion and geometric distortions, enabling higher nominal and effective spatial resolution. It is therefore anticipated that reconstructions using high resolution DW-SSFP dMRI data would offer advantages compared to tractography in lower resolution in vivo data.

Gains in anatomical detail are immediately apparent across states and resolutions in the orientation colour-coded DTI FA maps (Fig. 5a and Supplementary Fig. S13). The same animal has been imaged in vivo at (750 *μm*)^3^ and ex vivo at (400 *μm*)^3^ and (300 *μm*)^3^, and differences in fine structures can be appreciated in the cerebellum, brain stem and internal capsule. We further illustrate the benefits of pushing spatial resolution for in vivo acquisitions (Fig. 5b and Supplementary Fig. S14), where the same animal was imaged at (750 *μm*)^3^ and (580 *μm*)^3^ in vivo.

**Figure 5.**
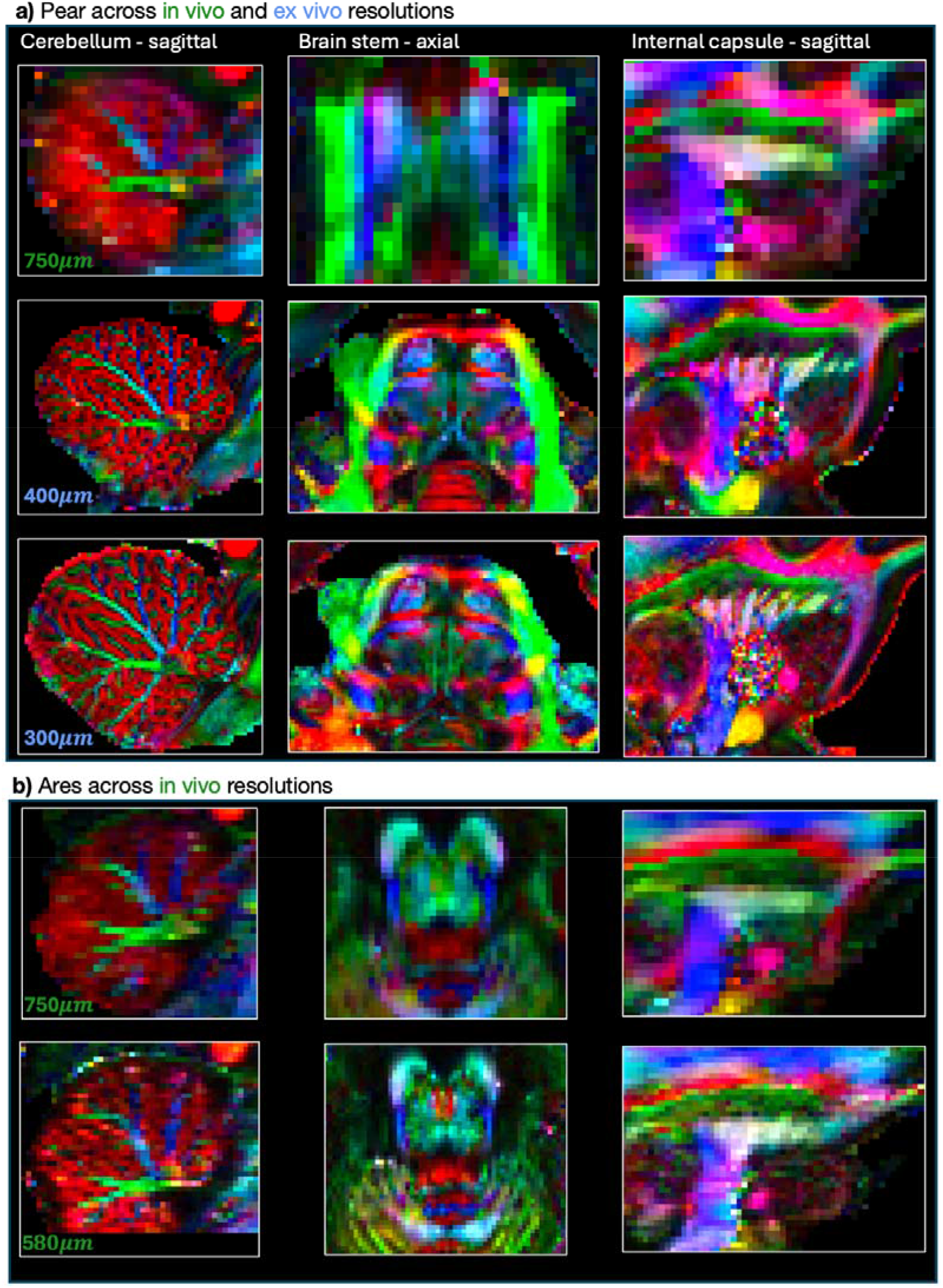
Comparison of zoomed in sections of the same brains imaged across resolutions and states exampled using colour-coded fractional anisotropy maps. a) NHP Pear was imaged in vivo (top row: 750) and ex vivo (middle row: 400, bottom row: 300). b) NHP Ares was imaged in vivo (top row: 750, bottom row 580). These data correspond to Ares (2) and Ares (3) described in Supplementary Table S2. Left column: zoomed sagittal slice of the cerebellum near the midline. Middle column: zoomed axial slice of the brain stem. Right column: zoomed sagittal slice of the internal capsule.

The downstream benefit from increasing spatial resolution in the data is also evident in tractography reconstructions of the relative locations of small, nearby and densely packed cortico-subcortical tracts^57^. Using in vivo and ex vivo data from the same animal, tractography provided good reconstructions for the striatal bundle (STB, external capsule) and extreme capsule (EmC) (Fig. 6a), with the former being medial to the latter in both low- and high-resolution data, but ex vivo tractography providing more focal reconstructions of the respective path density maps. The resolution benefit is more pronounced in the second example (Fig. 6b), with the relative location of the amygdalofugal bundle (AMF, anterior limb), uncinate fasciculus (UF), cingulum (CBP) and anterior commissure (AC) being more clearly delineated in the higher resolution data.

**Figure 6.**
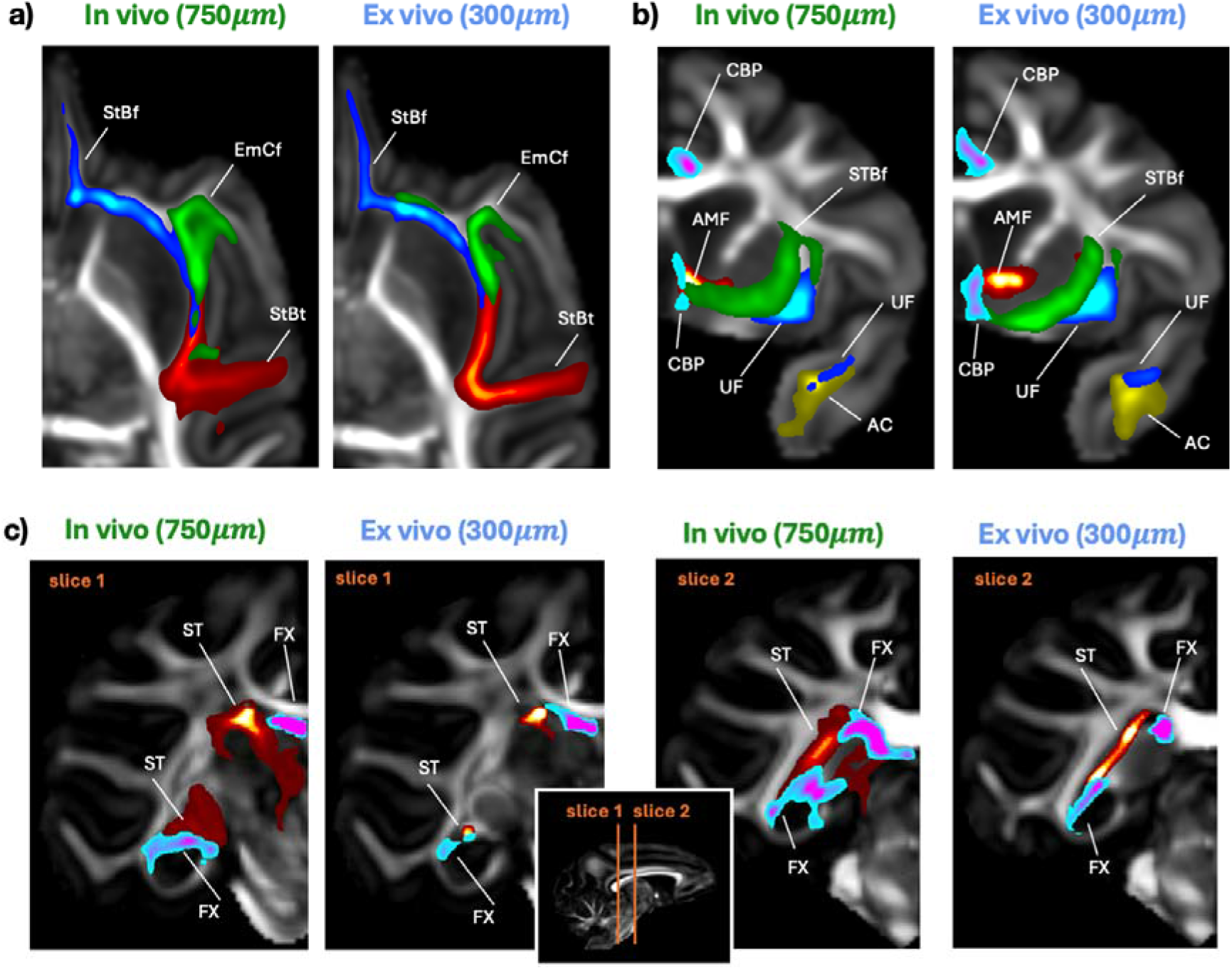
Advantages in tractography with enhanced spatial resolution. a-b) Single-animal tractography examples of white matter fibre bundles, reconstructed from in vivo and ex vivo data. Benefits from the higher resolution ex vivo data can be seen in preserving both localisation and relevant topographies of bundles that are running nearby to each other. c) Reconstructions of the stria terminalis (ST) and fornix (FX) bundles for the same brain in vivo and ex vivo showing gains in topography and bundle specificity across resolutions through coronal slices. AC: anterior commissure, AMF: amygdalofugal bundle, CBP: parahippocampal aspect of the cingulum bundle, StBf: striatal bundle/external capsule (frontal), StBt: striatal bundle/external capsule (temporal), EmCf: extreme capsule (frontal), FX: fornix, MB: Muratoff bundle, ST: Stria Terminalis, UF: uncinate fasciculus.

To further demonstrate the benefits of high-resolution diffusion imaging, we developed new tractography protocols for the stria terminalis (ST), which is particularly challenging as it runs in close proximity with the fornix. The two, however, connect different neighbouring limbic structures (amygdala and hippocampus respectively). Through an iterative protocol-development process using the (300 *μm*)^3^ ex vivo data and informed by histological literature^59,60^, we took advantage of the rich anatomical information, high spatial resolution and high signal contrast available in the ex vivo acquisition. The resulting protocols (detailed in Methods) enabled high-quality reconstructions of each bundle and were subsequently extended to the (750 *μm)*^*3*^ in vivo data of the same brain (Fig. 6c). Reconstructions of the fornix run nearby and medially to the stria terminalis and reach the hippocampus, while stria terminalis reaches the amygdala. Even if the pattern can be observed in both resolutions (suggesting generalisability of the proposed tractography protocol), the higher resolution ex vivo data yielded specific concentrated tractography reconstructions that did not propagate to nearby subcortical regions.

### Ultra-high-resolution mapping of whole-brain connectivity at 20

In addition to bundle-specific tractography, we performed probabilistic track-density imaging (TDI)^61^ to reconstruct whole-brain white matter pathways. This utilises the spatial continuity of tractography streamlines to super-resolve anatomical features by binning spatial distributions of streamlines at an arbitrarily high spatial resolution. Higher quality and spatial resolution data supports higher TDI resolutions (Supplementary Fig. S15 shows TDI differences between in vivo and ex vivo data across resolutions). In our case, starting from the (300 *μm*)^3^ ex vivo DW-SSFP data, we achieved more than an order of magnitude higher isotropic spatial resolution of 20 (Fig. 7 and Supplementary Videos 1-3). Major fibre bundles such as the corticospinal tract, corpus callosum and cerebellar peduncles can be seen in these contrasts (Fig. 7a). Fine white and grey matter structure can be similarly identified, for example, bifurcations and branch-like structure within the cerebellum (Fig. 7b,c), complex fibre architecture in the brain stem (Fig. 7d), the modular structure of the internal capsule (Fig. 7e), complex fibre crossings in the corona radiata and fibre fanning at the cortex (Fig. 7f,g). We anticipate that combining information from TDI and bundle reconstruction maps (Fig. 8) will enable unprecedented comparisons against whole-brain PS-OCT microscopy that will accompany these dMRI datasets, highlighting specific modes of success and failure for validating and improving tractography reconstructions.

**Figure 7.**
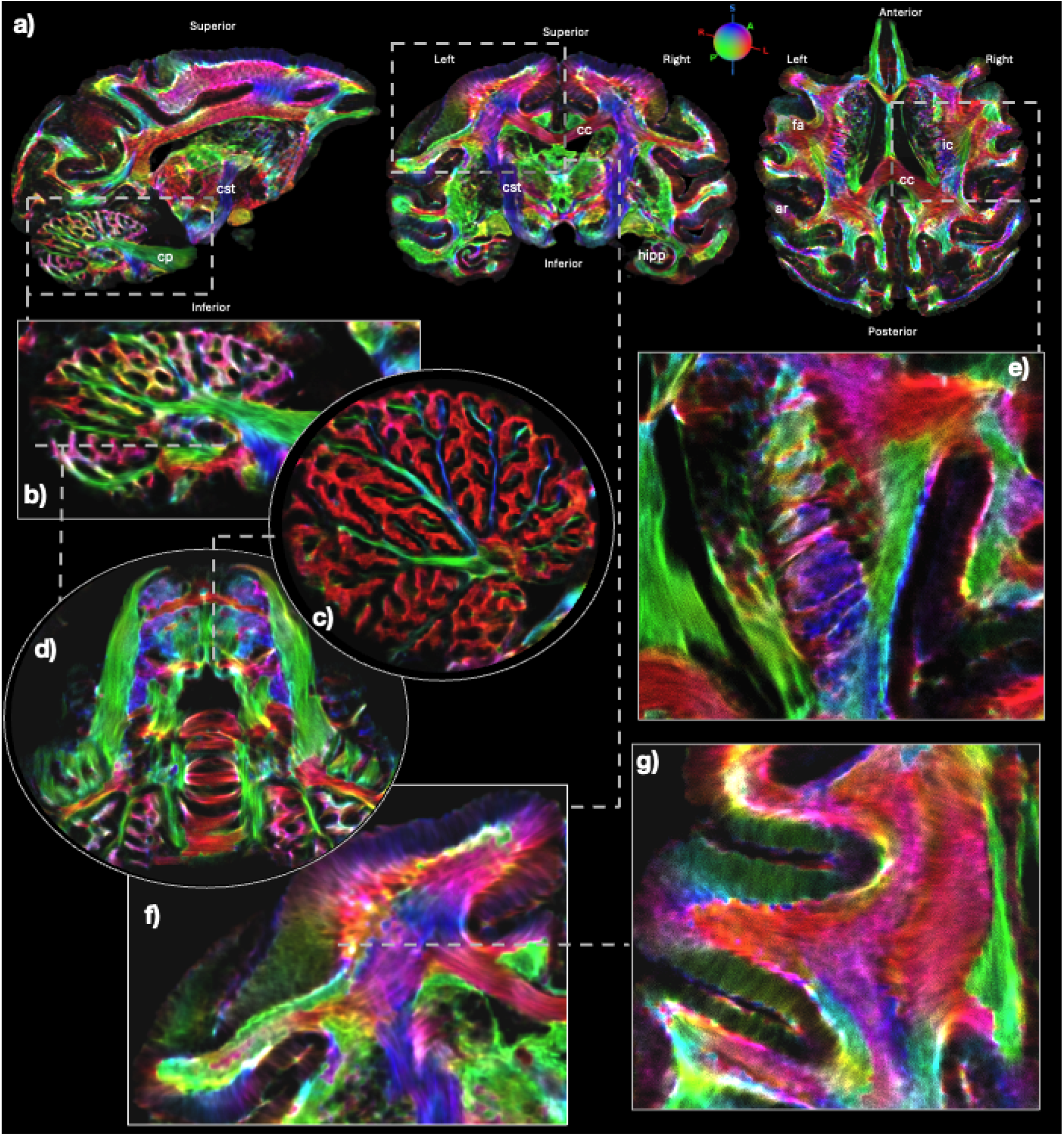
Whole brain ultra-high-resolution (20) ex vivo track density imaging (TDI). a) Whole-brain TDI images obtained from the ex vivo (300 μm)^3^ DW-SSFP data and colour-coded by the principal fibre DTI orientations and displayed across the three radiological axes with some major anatomical landmarks labelled. ar: acoustic radiation, atr: anterior thalamic radiation, cc: corpus callosum, cp: cerebellar peduncle, cst: cortico-spinal tract, ic: internal capsule, hipp: hippocampus. b,c,d) Magnified sagittal (b,c) and axial (d) views of the cerebellum highlighting the branch-like structure of white matter fibres. e) A magnified axial view of the internal capsule and surrounding region highlighting complex interweaving of white matter structures. f) A magnified coronal view of the corona radiata highlight complex fibre architecture and cortical fanning. g) A magnified axial view through the corona radiata highlighting cortical fanning.

**Figure 8.**
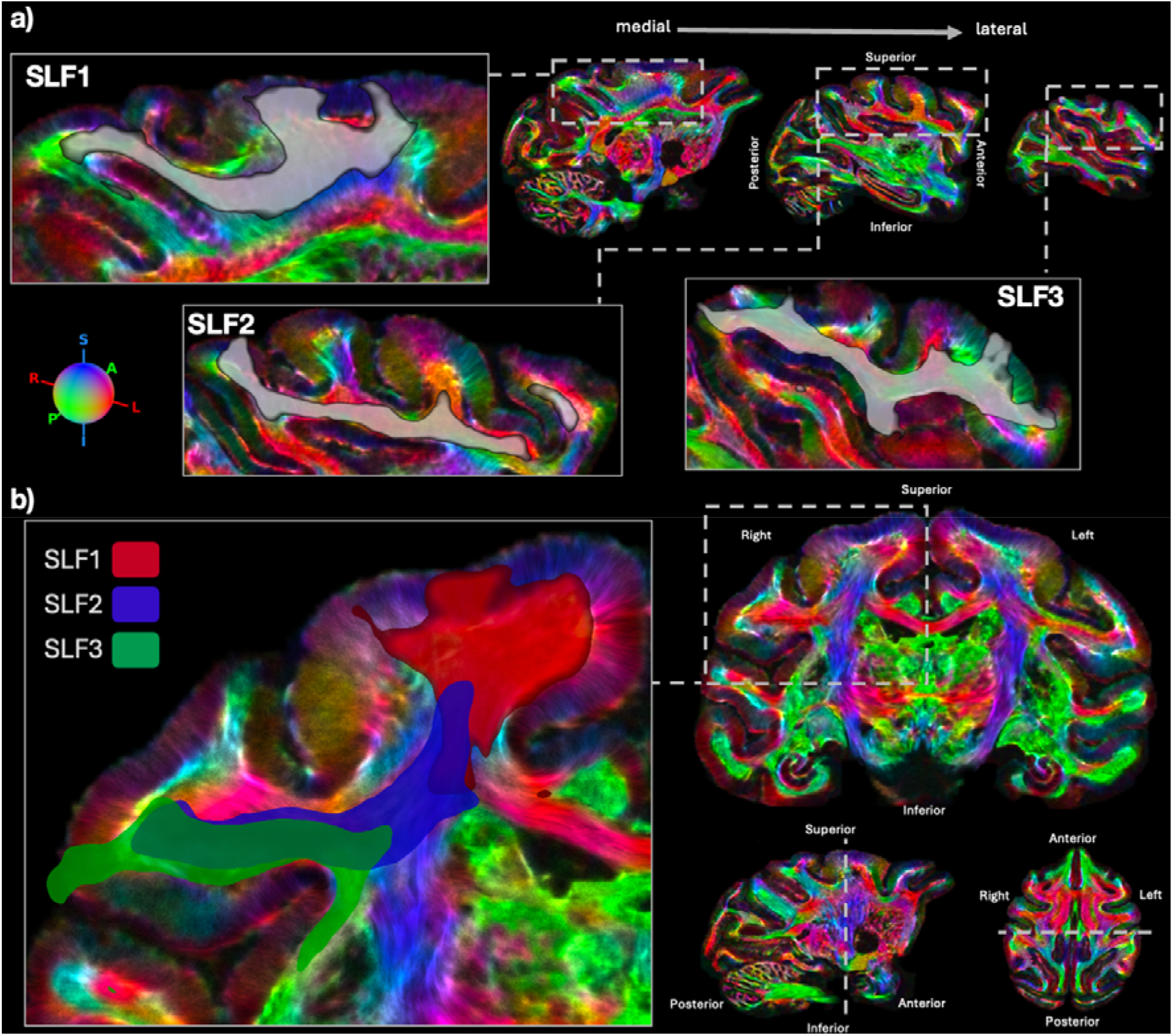
Imposing tractography-derived anatomical information to whole brain ultra-high-resolution ex vivo track density imaging (TDI). Tractography reconstructions of the three branches of the superior longitudinal fasciculi (SLF 1, 2 and 3), obtained from XTRACT NMT-space tract atlases, overlaid on TDI a) sagittal slices and b) a coronal slice of the tract density image. Note: to ease visualisation compute requirements, we use TDI at 30 here.

## Discussion

We have presented developments on acquisition and processing that enable high-resolution in vivo and ex vivo diffusion MRI of the NHP brain using a 10.5 T human-bore MRI scanner. By combining novel solutions to address challenges in scanning at UHF with optimisations of state-of-the-art approaches for acquiring and processing UHF data, we have demonstrated feasibility for high-quality NHP dMRI data and tractography across different tissue states and for isotropic resolutions up to 580 *μm* in vivo and 300 *μm* ex vivo, with b values up to 6,000 s/mm^2^. We have proposed a framework for UHF imaging of the same brains across resolutions and states, and developed end-to-end processing that spans image reconstruction and denoising, distortion correction, multi-modal registration, biophysical modelling and tractography. We have demonstrated gains in anatomical fidelity (Fig. 2f, 3c, 5, 6), enabled by advancements in acquisition and image processing (Fig. 1, 2, 3). We have shown how the higher resolution data can benefit tractography in localising and separating nearby subcortical bundles (Fig. 6) and in whole-brain ultra-high-resolution reconstructions (Fig. 7-8, Supplementary Fig. S15).

Resolving microscopic white matter architecture whilst maintaining whole-brain field of view in the living brain, remains an open challenge for imaging technologies and diffusion MRI is the only available technique to approach such competing demands. Our study opens new possibilities towards addressing this challenge on its own, but it is also a significant part of coordinated efforts for joint microscopy and MRI across whole brains within the Center for Mesoscale Connectomics (CMC). The data presented here are part of a wider dataset that further includes whole-brain PS-OCT and axonal tracing of the same brains and will additionally acquire both in vivo and ex vivo dMRI and PS-OCT for the human brain. Advances in imaging resolution made through the presented work have pushed our ex vivo dMRI imaging close to the 200 *μm* section thickness of PS-OCT performed in the same brains^62^. These data will be subsequently used to jointly model brain microstructure and white matter architecture across scales, modalities and species to transfer knowledge and methods towards in vivo imaging. Recent studies using multimodal, multiscale and multistate (in vivo and ex vivo in the same brain) approaches have driven advancements in this direction and our understanding of neuroanatomy. For example, the fusion of multi-contrast MRI and microscopy of the same NHP brain has been used to refine connectivity estimates^23,24,63^, taking advantage of the unique, yet complementary, information encoded in each modality. Similarly, multimodal imaging has been used to compare and validate estimates of tractography in the developing human brain^64^. However, multimodal, multiscale and multistate imaging of the same brain and for multiple brains is a complex task that has not yet been fully tackled, which the CMC addresses.

A number of prior studies have reported NHP data with sub-millimetre resolutions either in vivo^27,65– 68^ or ex vivo^23,28,28,29,32,51,69–75^ (Supplementary Tables S5 and S6). All previous ex vivo studies have used high-gradient preclinical systems that due to smaller bore size have less challenging RF/field/gradient inhomogeneities, but cannot be used to scan NHPs in vivo. Our achieved resolutions at 10.5 T are better or comparable to previously reported ones with high-gradient preclinical systems, with the exception of Bihan-Poudec et al.^27^ and Tounekti et al.^66^ that achieved 400 *μm* and 500 *μm* isotropic resolutions in vivo, respectively, using human 3 T scanners, but at a low b value and a few volumes. A number of aspects make our work unique, advancing prior studies: i) No previous work has used the same setup for pushing resolution across tissue states of the same brain. Imaging in vivo and ex vivo is enabled using a large bore 10.5 T scanner, but represent quite different set of challenges, in dealing with distortions, signal uniformity, image reconstruction and contrast. Addressing these challenges enables direct cross-state comparisons whilst minimising scanner differences, and provides the foundation for integrating diffusion MRI with tract tracing and microscopy (e.g. PS-OCT) in the same brains (that are developed in parallel in our consortium^25,62,76^), allowing validation and refinement of tractography and microstructure models. ii) Achieving the presented developments using a human UHF scanner makes them in principle more directly transferable to human in vivo (DW SE-EPI) and ex vivo (DW-SSFP) UHF 10.5 T imaging. Lessons learnt on coil and gradient performance^38^, pulse sequence behaviour, signal and noise uniformity/stationarity, modelling assumptions, tolerance to geometric distortions and ability to correct, dMRI reconstruction and processing in such a prototype UHF system can be translated to human UHF imaging advances. For instance, we identified that higher order eddy current models offer better distortion correction performance for this data and this is likely to hold for human dMRI processing; or that single-shot SE EPI with moderate in-plane acceleration can deem high-resolution in vivo UHF data with addressable susceptibility-induced distortions. In fact ongoing developments for dMRI imaging of living humans at 10.5 T, building upon this work, have led to preliminary high-res DW SE-EPI human data^30^ at 10.5 T.

Previous studies using DW-SSFP have been performed at lower field strengths, including ex vivo human brain imaging at 3 T^77^ and 7 T^37,78^ achieving sub-millimetre resolutions, with a recent 7 T approach reaching an isotropic spatial resolution of 850 *μm*^*78*^. For the macaque, 3 T ex vivo DW-SSFP acquisitions^77^ achieved submillimetre (700 *μm*) in plane but with larger slice thickness profiles (1.4 mm), which is substantially lower than the DW-SSFP resolution reported here. In addition, SNR efficiency calculations using ex vivo relaxometry parameters at 10.5 T^31^ have demonstrated increased SNR performance over alternative sequences (such as diffusion-weighted spin echo or stimulated echo) at high field strengths^31,79^.

In this work, we utilised a prototype human scanner with ultra-high field strength (10.5 T, the second strongest field strength human MRI system in the world), but relatively moderate gradient strength (70 mT/m). Modern ultra-high performance gradient systems offer benefits for diffusion imaging both at high field strengths^16,20^ and also at conventional field strengths^10,21^, because they lead to higher SNR/CNR by shortening the echo time (TE). However, SNR is determined not only by TE but also by the intrinsic SNR available at TE=0, i.e. intrinsic SNR available at a given magnetic field, which is strongly field-dependent and benefits high magnetic fields. Of course, the most ideal situation is combining ultra-high magnetic fields with ultra-high-performance gradients. This combination is currently not available in the 10.5 T system, but new developments in the near future will move towards this direction (the Impulse gradient insert by Siemens is already planned for the 10.5 T scanner). All of the RF coils developed in this project were designed to fit in such a gradient set in anticipation of this development. Despite the current challenges associated with high magnetic fields, like 10.5 T, we demonstrated in this work that we can push resolution and contrast (achieving relatively high b values) for both in vivo and ex vivo brain tissue states, even in the absence of ultra-high gradients, utilising the field dependent gains in the intrinsic SNR. These results can also be taken as projecting future gains that will be available when the Impulse gradient is incorporated at 10.5 T.

The resolutions achieved here for the macaque brain are on par with what would be considered high-resolution for the human brain. The macaque brain volume is roughly 15 times smaller than the human brain^68,80^. Assuming roughly similar geometric complexities, an isotropic resolution scaling factor of 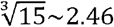 could be used as a guide for identifying similar voxel volume/brain volume ratios. Our in-vivo NHP resolutions of 580 *μm* and 750 *μm* would therefore be on par with human in vivo data of 1.4-1.8 *mm* isotropic resolution (i.e. HCP Lifespan remit^81^), while the ex vivo 300 and 400 *μm* would correspond to sub-millimetre human ex vivo 700-950 *μm* isotropic data at the b values acquired, with 850 *μm* amongst the highest ex vivo resolution for human brains acquired at 7 T^78^. Notably, the resolutions achieved here surpass the recommendations of the NHP neuroimaging and anatomy project and even approach the 500 *μm* guide for anatomical macaque imaging^68,80^, which seeks to maintain an isotropic resolution of half the cortical thickness. With our current spatial resolutions, we managed to reconstruct a range of white matter bundles and identify fine anatomical features, as exampled in Figure 6.

In summary, we have presented a framework for high resolution diffusion MR imaging of the in vivo and ex vivo NHP brain using an ultra-high field strength human-bore scanner. Our acquisitions have been designed and optimised to a degree that allow using established diffusion MRI preprocessing tools where possible, whilst also providing novel solutions and end-to-end processing pipelines, from image reconstruction to tractography, for challenges specific to ultra-high-field and ex vivo imaging. The current data, in addition to upcoming data acquisitions, along with all processing pipelines, containerised workflows, and reusable web services, are openly shared to support reproducibility and future integration with whole-brain microscopy^3,4^.

## Online Methods

### Hardware & Coils

All data were acquired using the MAGNETOM (Siemens, Erlangen) 10.5 T scanner at the CMRR, University of Minnesota. The scanner has a bore size of 88 cm (60 cm accessible bore within gradient insert) and is fitted with SC72D gradients (70 mT/m, 200 T/m/s slew rate). Both in vivo and ex vivo data were acquired using custom-made (SNR optimised and receive/transmit signal homogenised) coils. RF coils were developed through the duration of these data (see Supplementary Fig. S1 for examples) and the final in vivo coil has 8-channel/32-channel transmit/receive arrays, while the ex vivo coil consists of 8-channel/40-channel transmit/receive^82^.

### Animal preparation

Six macaques were used in the study so far (five male rhesus and one female cynomolgus, age range 4-9 years old). For in vivo acquisitions, on scanning days, anesthesia was first induced by intramuscular injection of atropine (0.5 mg/kg), ketamine hydrochloride (7.5 mg/kg), and dexmedetomidine (13 µg/kg). Initial anesthesia was maintained using 1.0%–2% isoflurane mixed with oxygen. Each animal was wrapped in warm packs to maintain body temperature. A circulating water bath was used to provide additional heat. A ventilator was used to prevent atelectasis of the lungs, and to regulate CO_2_ levels. The animals were observed continuously, with vital signs and depth of anesthesia monitored and recorded at 15-min intervals. Rectal temperature (~37.6 °C), respiration (10–15 breaths/min), end-tidal CO_2_ (25–40), electrocardiogram (70–150 bpm), and oxygen saturation (>90%) were monitored using an MRI compatible monitor (IRADIMED 3880 MRI Monitor, Orlando, FL, USA).

After in vivo scanning, animals underwent surgery with AAV injection for axonal and neuronal viral labelling. Animals each received 6-10 microinjections of AAV delivered to frontal and parietal targets. Two months after surgery, animals were euthanized using pentobarbital following ketamine/dexmedetomidine sedation and transcardially perfused using phosphate-buffered saline (PBS) followed by 4% hydrogel monomer solution (40% w/v acrylamide, 2% w/v bisacrylamide, 10X PBS, 8% w/v paraformaldehyde and distilled water). Brains were extracted and kept in 4% hydrogel monomer solution (HMS) for one week. Following, brains were kept in PBS until imaged^83^. Prior to ex vivo imaging, brains were brought to room temperature and transferred into the form fitting shells and filled with a susceptibility matched electronic liquid (3M Flourinert FC-3283). Small air bubbles were removed using rotational tilting as well as using a soft catheter tip. Brains were left to sit overnight to alleviate any recurrence of air bubbles previously undetected and then transferred to the magnet room in the morning of the scan. After ex vivo MR data acquisition, the specimens were transferred back into PBS and stored in the refrigerator.

### Ethics

Experimental procedures were carried out in accordance with the University of Minnesota Institutional Animal Care and Use Committee and the National Institute of Health standards for the care and use of NHPs. All subjects were fed ad libitum within a light and temperature-controlled colony room. Animals had access to ad lib water.

### Acquisition protocol development

Given that the 10.5 T human scanner is a prototype system, there is limited knowledge on efficient acquisition and dMRI protocols^14^. We therefore relied on an iterative procedure for optimised acquisition development that considered and tested the various trade-offs, from image reconstruction, distortion handling and quality assessment all the way to tractography, with outcomes feeding back and informing next steps.

Both in vivo and ex vivo acquisitions have unique advantages and challenges which must be considered, whilst pushing both spatial and angular resolution. For in vivo acquisitions (inherently more limited by acquisition time and affected by potential sample motion), we opted for diffusion-weighted (pulsed-gradient) spin-echo 2D echo planar imaging (DW SE-EPI), as DW SE-EPI allowed very high quality human dMRI data at 7 T for the Human Connectome Project^16^, still considered state of the art, and we anticipate developments to be similarly applicable to in vivo human dMRI at 10.5 T. To mitigate for the higher susceptibility-induced distortions at ultra-high field, which can be severe for EPI data, we assessed high in-plane acceleration factors (GRAPPA factors up to 4), in combination with iterative B_0_ shimming and power calibrations for each animal using a transmit B_1_ fieldmap. We also used blip-reversed *b=0* acquisitions^45^ to estimate an off-resonance fieldmaps and explored different phase encoding directions (AP/PA vs LR/RL). We tested different spatial resolutions including 750 *µm* and 580 *µm* and, through the optimisation of acquisition hardware, protocols and image processing, we have now robustly achieved an isotropic spatial resolution of 580 *µm*. At the same time, improvements in hardware, including scanner gradients and coils, have enabled more homogeneous signal coverage and manageable geometric distortions (Supplementary Fig. S1). For q-space sampling, we relied on the multi-shell protocol published before^14^, as providing good compromise between diffusion contrast, angular resolution and SNR.

For ex vivo acquisitions, we opted for diffusion-weighted steady-state free precession (DW-SSFP)^84– 86^, which has been shown to provide advantages in SNR efficiency for ex vivo imaging in comparison to spin-echo and stimulated echo sequences^56,77,79,87^, alongside improved SNR-efficiency at higher magnetic field strengths^37^. As DW-SSFP is a 3D technique with a short TR, a single-line k-space readout was incorporated whilst keeping acquisition times reasonable, and hence acquired imaging volumes are effectively distortion-free. We also explored spatial resolutions (500, 400 and 300 *µm*) and b-values (up to ~8000 s/mm^2^), with the chosen multi-shell scheme providing the best compromise between angular contrast, sensitivity to fibre crossings and signal to noise ratio (Supplementary Fig. S2).

In vivo 2D DW SE-EPI acquisitions on anesthetized macaques used an isotropic spatial resolution of 750 *µm* or 580 *µm* with two b-value shells (b=1000, 2000 s/mm^2^) and 54 volumes per shell. Four repeats for each of the two phase-encoding (PE) directions anterior-posterior (AP)/posterior-anterior (PA) were acquired, allowing for blip-reversed informed correction of susceptibility-induced distortions. We used a parallel imaging (GRAPPA) acceleration factor of three, and a 75% partial Fourier factor. The number of slices acquired for each brain were in the 70-80 slices range to accommodate for the different brain sizes, whilst ensuring whole brain coverage. Hence, repetition times varied slightly across acquisitions (around 8.5s for 750 *µm* and 9.2s for 580 *µm*, Table 1 and Supplementary Table S2). Echo time was 66ms and 79ms for the 750 and 580 *µm* resolution, respectively. Anatomical images were obtained with a T2-weighted Turbo Spin Echo (TSE) sequence (TR⍰1=⍰13000⍰1ms, TE⍰1=⍰1100⍰1ms, flip anglel1=l190°, voxel size = 500 *µm* iso, matrix sizel1=l1260×320, GRAPPA⍰1=⍰12, bandwidth⍰1=⍰1504⍰1Hz/pixel). In vivo data were collected from five NHPs (Curly, Larry, Pear, Ares, Gandalf).

Ex vivo 3D DW-SSFP acquisitions used an isotropic spatial resolution of 400 *µm* or 300 *µm* with two shells (b_eff_ ~3200, 5700 s/mm^2^) and 60 volumes per shell (Table 1). Protocol parameters were optimised for each animal based on T1 and T2 values of the fixed tissue and are provided in Supplementary Table S3 for each of the imaging sessions so far. Inherently, the b-value is not directly applicable to DW-SSFP acquisitions. However, to facilitate interpretation and to ease interaction with standard image processing tools, we calculate and report effective b-values which are derived from the estimated diffusion attenuation and an approximate diffusion coefficient (d_ex-vivo_ ~ 2·10^-4^ mm^2^/s)^56^. DW-SSFP acquisition time per volume was approximately 15 minutes for 400 *µm* and 27 minutes for 300 *µm*, with a total ex vivo scan time of approximately 28 hours and 54 hours respectively. Anatomical T2-weighted TSE images were acquired with the following parameters (TR = 3000 ms, TE = 250 ms, flip angle = 90°, voxel size = 120 *µm* iso, matrix size = 492×800, slices = 416, GRAPPA = 3, bandwidth = 202 Hz/pixel). Ex vivo data have been collected so far from two NHPs (Moe, Pear).

Diffusion attenuation in DW-SSFP is dependent on tissue relaxation (T1 and T2) properties and several sequence parameters (TR, flip angle, diffusion gradient properties)^54^, which consequently impose a dependency on field inhomogeneities (via the B1 field). As such, we also acquired quantitative voxel-wise T1 and T2 relaxation maps for the samples and B1 field maps. T1 mapping was performed using a Magnetization Prepared 2 Rapid Acquisition Gradient Echoes (MP2RAGE) sequence with an isotropic spatial resolution of 1 mm. T2 maps were reconstructed from multi-echo spin-echo data, acquired at 875 *μm* by 875 *μm* by 1 mm resolution, using a custom extended-phase graphs (EPG) fitting routine^49^. B1 field maps were scanner-reconstructed from turbo FLASH data acquired using a 1.3 by 1.3 mm matrix and a 2 mm slice thickness. T1, T2 and B1 maps were linearly registered to the DW-SSFP data for subsequent modelling. Full details on parameter mapping can be found elsewhere^31^.

For both in vivo and ex vivo acquisitions, complex (magnitude and phase) data were stored for the diffusion acquisitions.

### Image reconstruction and denoising

Data from different channels were combined using a sensitivity encoding (SENSE1) reconstruction^39^ to minimise noise floor in the combined data. SENSE1 reconstruction of the DW SE-EPI data was performed on the scanner for in vivo data using the CMRR multiband C2P package^88^ (http://cmrr.umn.edu/multiband/). For the ex vivo DW-SSFP data, the acquisition sessions had to be segmented into around 30 partitions to support reasonable data files, which also only allowed for Adaptive Combine coil-combination to be used for the online reconstruction. We therefore performed offline reconstruction for ex vivo data, as Adaptive Combine leads to higher noise floor than SENSE1 coil combination^39^ (Supplementary Fig. S4). Data were subsequently denoised in the complex domain using NORDIC^40^ (Supplementary Fig. S5), as complex denoising has been shown to reduce both noise-induced variance and noise floor, improving both accuracy and precision of subsequent derived features^41^.

### Processing

We developed end-to-end processing pipelines for in vivo and ex vivo data, generalisable to NHP (macaque) and human data. Further, the ex vivo pipeline also generalises across acquisition types, including DW-SSFP and conventional (single-line readout) DW-SE acquisitions. To aid portability and reproducibility, pipelines are also available through dockerised containers. Conceptually, the pipelines include denoising, distortion correction steps for in vivo data, quality control, diffusion tensor modelling and fibre orientation distribution estimation, standard space registration and WM bundle tractography, all outlined in detail below. Both pipelines are available as Docker containers, are GPU-enabled (but also have the option to run without GPUs), and allow flexible switching on/off of pipeline steps.

#### Gibbs unringing and bias field correction

For both in vivo and ex vivo, we corrected for ringing artefacts using the Gibbs (2D for in vivo, 3D for ex vivo) unringing function implemented in MRtrix (v3.0.7)^42–44^ (Supplementary Fig. S6). This was performed after denoising to avoid corruption of signal property assumptions during denoising but prior to any further corrections. For the ex vivo data, we additionally applied Gibbs unringing to T1 and T2 parameter maps prior to relaxation mapping. For the in vivo data, following Gibbs unringing, we corrected B1 inhomogeneities using the N4 ANTs algorithm implemented in MRtrix (v3.0.7)^44^. We did not perform this step ex vivo, instead B1 inhomogeneity maps were used during diffusion modelling.

#### Signal intensity drift correction

As temporal scanner instability, and potentially other factors such as changes in tissue temperature, may introduce changes in signal intensity^89^, we implemented signal drift corrections, or intensity normalisations, for both the in vivo and ex vivo data (Supplementary Fig. S7). We followed the general approach implemented in the Human Connectome Project (HCP) pipelines^8,90^. Specifically, dMRI volumes were split into subsets and the mean b=0 signal intensity was calculated across all b=0s within a given subset. The first subset was used as a baseline against which all subsequent ones were normalised against, using the ratio between the baseline mean b=0 intensity and the mean b=0 intensity of each subsequent subset. For in vivo, we considered subsets of ~100 volumes (i.e. both shells for each phase-encode direction and repeat). For ex vivo, due to the longer scan time, we considered subsets of ~20 volumes. Additionally, we restricted the calculation of the mean b=0 intensities to the brain (excluding ventricles) using template-space masks non-linearly aligned to the native diffusion space. This was necessary to avoid contamination from any residual PBS in the ventricles.

#### Susceptibility, motion and eddy current distortion correction

In vivo data were motion, susceptibility distortion and eddy current distortion corrected, including all these corrections in a single interpolation step, using FSL-topup and FSL-eddy^45,46^.

Eddy currents, arising from rapid diffusion gradient switching, are typically modelled as linear or quadratic spatial fields. Whilst quadratic modelling is sufficient for 3 T or even 7 T data^46^, we evaluated the performance of linear, quadratic and cubic eddy current modelling for our ultra-high field strength data. Eddy current modelling parameters were evaluated by independently correcting reverse phase-encoding pairs (AP/PA) and correlating the estimated parameters across volumes. A good model should yield high AP–PA correlations in the estimated parameters, while low parameter correlation across independent runs of the same data suggest overfitting or inability to capture more complex eddy current effects. In addition to identifying the best spatial model for capturing eddy current off-resonance fields (Supplementary Fig. S8), we explored smoothing kernels in FSL-eddy iterations, as a way to increase robustness to large displacements and/or positional shifts during the acquisition (FWHM: 10, 5, and 0 mm smoothing kernels) (Supplementary Fig. S9). Finally, we used automatic outlier detection and replacement^46^, estimated and corrected the susceptibility and sample motion interactions and configured eddy optimally for ultra-high field distortion corrections.

For DW-SSFP data, which are distortion free, none of the above corrections were necessary. Instead, a rigid-body transform of all volumes to the first was used to account for spatial drifts over the long acquisition.

#### Quality Control

We calculated image quality metrics including the signal to noise ratio (SNR) of the b=0 volumes and angular contrast to noise ratio (CNR) of each shell, using FSL-eddyqc^91^. This was applied for both in vivo and ex vivo data.

#### Skull stripping

In vivo, skull stripping was performed using FSL’s bet4animal^68,92^ on the anatomical T2w image and the resultant brain mask registered to the native dMRI space. Ex vivo, brain masks were generated through voxel intensity thresholding, clustering and erosion/dilation. The pipelines perform skull stripping following these procedures by default, however, users may instead provide a pre-computed brain mask.

#### Diffusion tensor estimation

For both in vivo and ex vivo, we performed diffusion tensor modelling using the lower shell of the data (b=1000 s/mm^2^ for in vivo, b_eff_ ~ 3,000 s/mm^2^ for ex vivo), exampled in Supplementary Fig. S10. In vivo, this was performed using the standard diffusion tensor model. Ex vivo, we used a DW-SSFP modified diffusion tensor model^54,78^ and an MCMC estimation implemented in CUDIMOT^48^ to account for the specific signal properties of DW-SSFP data and their dependency on tissue parameters (T1 and T2 tissue properties) and B1 field inhomogeneities.

### Registration and multi-modal template creation

In order to compare brains in standard template space, and to facilitate standard space landmark-based tractography, we registered both in vivo and ex vivo data to the standard NIMH Macaque Template (NMT v2)^53^ macaque space, which includes high resolution (500 *µm* isotropic) T1-weighted templates. We explored two approaches for registration. Firstly, a conventional approach of non-linear registration^93^ using the native space fractional anisotropy (FA) to the T1w NMT template. Registering FA to T1w template is common practice for NHP data, particularly when anatomical scans are not available. Secondly, we explored the recently developed FSL-MMORF^47^ multi-modal non-linear registration framework, which allows transformations to be informed by multimodal features, including the diffusion tensor and principal diffusion directions. To achieve this, we developed NMT-space diffusion (FA and tensor) templates through a multi-step template development pipeline, described below, using a previous external dataset consisting of six ex vivo macaque brain scans^51,69^ available through PRIME-DE^94^: DW-SEMS, TE/TR: 25 ms/10 s; matrix size: 128 × 128; resolution: 600 *µm* isotropic; number of slices: 128, with 16 (b = 0 s/mm^2^) and 128 diffusion-weighted (b = 4000 s/mm^2^) volumes.

Specifically, for template creation, each of the six subjects’ FA maps were nonlinearly registered to NMT-space using FSL-FNIRT before constructing a cohort template using the ANTs build template tool^95,96^ with five refinement iterations. Separately, native space tensor data were warped to NMT space using the FNIRT derived transforms, including re-orientation of tensor elements^97^, providing an initial NMT-space tensor target. The “refined” affine and nonlinear transforms from the ANTs build template step were combined and applied to native space tensors with the FNIRT NMT-space tensors from the previous step as targets, resulting in a set of NMT-space tensors. These were subsequently averaged to generate the NMT-space tensor template and decomposed to extract standard diffusion tensor metrics, including FA, MD, eigenvalues (L1, L2, L3) and eigenvectors (V1, V2, V3). Once the NMT-space multimodal template was established, we used MMORF to estimate native space to NMT-space registration fields for our data, using the FA and tensor maps to inform registrations.

The pipelines include both registration options (unimodal or multimodal), which the user may select in the configuration file. Further, for human data, the pipeline automatically updates to the MNI152 standard space and uses the FSL_HCP1065 standard space atlases available within FSL for multimodal MMORF registrations.

### Fibre orientation mapping

For estimation of fibre orientation distributions (FODs), we explored both parametric and non-parametric ways for spherical deconvolution, as the former showcases feasibility for further microstructural modelling using both the in vivo DW SE-EPI and ex vivo DW-SSFP data. For the DW SE-EPI data, we considered the multi-shell ball & sticks model^98^, estimating using MCMC up to three fibre orientations per voxel, deconvolving the signal against the conventional DW-SE stick response kernel available in FSL-bedpostX. For the DW-SSFP data, we first explored the standard DW-SE ball & sticks with an effective b-value approximation^56^. Whilst this provided reasonable estimates (Supplementary Fig. S12), it had lower sensitivity for multiway crossings, as the DW-SSFP deconvolution kernel is different (“fatter”) than the conventional DW-SE kernel^77^. Therefore, we built upon previous work and developed a DW-SSFP extension of the ball & sticks model, considering up to 3 orientations per voxel and signal dependences on T1, T2 and B1 maps (similar to how the DW-SSFP tensor model was fit). We estimated this model using a GPU-based MCMC approach implemented in CUDIMOT^48^.

In addition, multi-tissue multi-shell (MSMT) constrained spherical deconvolution (CSD) was performed on the DW-SSFP data using MRtrix3^44^. To do so, we also relied on an effective b-value approximation^56^ to feed into CSD, as diffusion-weighting in DW-SSFP does not correspond to discrete well-defined b-values^54^. Tissue-specific response functions for white matter, grey matter, and cerebrospinal fluid were estimated in an unsupervised manner from the data^99^. Fibre orientation distributions (FODs) were then computed for each voxel using the three-tissue response functions.

By default, the ex vivo pipeline operates in DW-SSFP mode, however, the user may select the DW-SE mode if they have acquired conventional single-line readout DW-SE data. In this mode, the ex vivo pipeline uses the standard DW-SE models for diffusion tensor and crossing fibre modelling.

### Tractography

Probabilistic tractography using landmark-based WM bundle protocols defined in NMT space was performed using FSL-XTRACT^50,51^, allowing for the reconstruction of 60 major white matter fibre bundles. These include association, commissural, projection and limbic cortico-cortical fibre bundles, as well as cortical-subcortical bundles, such as the cortico-striatal, amygdalofugal and Muratoff bundles^57^. A full list of reconstructed tracts is provided in Supplementary Table S4. Default probabilistic tractography termination criteria were used (curvature threshold: ±80°, max streamline steps: 2000, subsidiary fibre volume threshold: 1%, randomly sampled initial fibres in case of fibre crossings in a seed location, no minimum length constraint, loop-checking and termination) with a step size of 150 *µm*. Notably, these tractography reconstructions have by design one-to-one equivalents to corresponding human reconstructions, to facilitate future comparative neuroanatomy studies.

We further developed new tractography protocols to reconstruct and resolve the fornix (FX) and stria terminalis (ST). These protocols used a seed-exclusion approach where a single seed mask is placed in the main body of the respective tract and exclusions used to restrict streamlines. For both bundles, a bounding exclusion box was placed superiorly, posteriorly and anteriorly to the bodies of the tracts, plus a midline exclusion to prevent leakage into the contralateral hemisphere. Exclusion masks were also placed in the corpus callosum, internal capsule, optic tract, anterior commissure, amygdala-fugal pathway and in the white matter of the temporo-parietal junction. For the ST, an additional exclusion was placed in the body of the FX.

In addition to bundle-specific tractography, we performed whole-brain tractography and reconstructed spatial histograms of the streamline density, known as track density images (TDI)^61^. Probabilistic tractography^55^ was performed by seeding randomly throughout the whole-brain whilst restricting the total number of steps to 300, with a step size of 10 *µm*, 50 samples per seed location with approximately 320 million randomly distributed seed voxels and binning the resultant spatial distribution of streamlines into 20 *µm* voxels using the ex vivo data (300 *µm* native resolution), thus generating an ultra-high resolution whole-brain track density image. Due to the high memory requirements (peak memory required: 545 GB) of 20 *µm* TDI, processing was performed using the CPU implementation of FSL probtrackx2 and took approximately 13 days of compute time. The 20 *µm* TDI results were visualised, and Supplementary Videos 1,2,3 were created, using custom python-based visualisation software. Supplementary Fig. S15 shows indicative TDI maps at different binning resolutions, for in vivo and ex vivo data, highlighting significant gains in anatomical detail with increased native imaging resolution (i.e. ex vivo compared to in vivo) and convergence in detail when comparing across TDI resolutions for the same data.

## Data Availability

Raw and minimally processed NifTI files for the six in vivo and three ex vivo datasets presented are available on brainlife.io: https://doi.org/10.25663/brainlife.pub.62.

## Code Availability

Pipelines are available to download and install through GitHub (https://github.com/SPMIC-UoN/CMC_invivo_dMRI_pipeline, https://github.com/SPMIC-UoN/CMC_exvivo_dMRI_pipeline). Without a container, pipeline external requirements include FSL (v6.0.7.18+), MRtrix3 (v3.0.7+) and NORDIC (https://github.com/SteenMoeller/NORDIC_Raw, and the EDDEN wrapper https://github.com/SPMIC-UoN/EDDEN). Additionally, the ex vivo pipeline requires the CUDIMOT (including the DTI and crossing fibres models, https://github.com/SPMIC-UoN/cudimot) when operating in DW-SSFP mode, and the 3D Gibbs unringing algorithm (https://github.com/jdtournier/mrdegibbs3D). Note: CUDIMOT modelling requires a CUDA toolkit and a GPU. Docker containers do not support image denoising due to licencing restrictions.

## Acknowledgements

This work is funded by NIH grant UM1NS132207 and the Centre for Mesoscale Connectomics. SW and SNS acknowledge funding from the European Research Council (ERC Consolidator — 101000969 to SNS). BCT is funded by a Sir Henry Wellcome Postdoctoral Fellowship (Wellcome Trust) [222829/Z/21/Z]. KLM is funded by a Wellcome Trust Senior Research Fellowship [224573/Z/21/Z]. The work is also supported by NIH grants P41EB027061 (JZ, KU, GA), R01EB031765 (JZ) and P30DA048742 (JZ, SH) and ERC grant 101220843 (UltraDeepMood to DF).

## Notes

### Competing Interest Statement

The authors have declared no competing interest.

### Summary of Updates

Manuscript now includes higher resolution dMRI for both in vivo and ex vivo imaging, and illustrates additional gains from enhanced spatial resolution.

