## Supplementary Materials for "10.5 Tesla High-Resolution Macaque Brain MRI for In vivo and Ex vivo Connectivity Studies"

#### *Hardware upgrades*

Acquisition of dMRI data at UHF presents challenges in geometric distortions (in vivo), RF/field/gradient inhomogeneities and signal coverage. We trialled multiple acquisition strategies to optimise these factors whilst maintaining useful signal/contrast to noise ratios and optimising spatial and angular resolution. In addition to acquisition protocols, acquisition hardware was also developed and optimised[1,2]. Supplementary Fig. S1 demonstrates examples of data using older versions of gradients and head coils, demonstrating how inhomogeneous signal coverage across the whole brain (yellow arrows) has been resolved with the hardware upgrades.

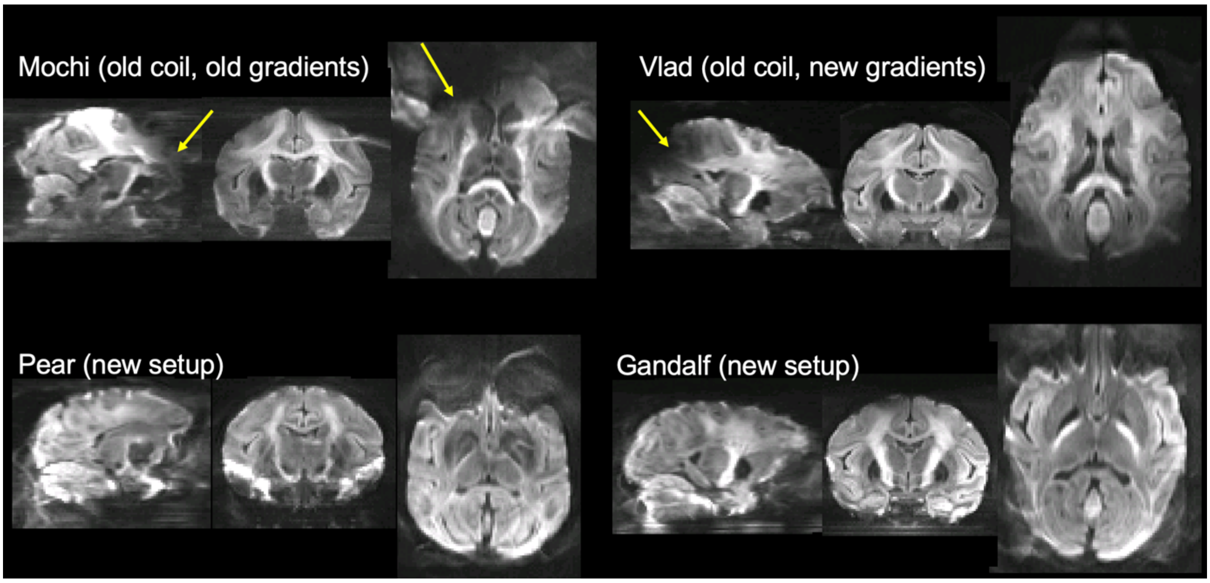


***Supplementary Fig. S1*** *- Iterations of in vivo data acquisition showing improvements in data signal homogeneity and geometric distortions with newly developed RF and gradient coils. Arrows indicate regions of signal dropout.*

### *Q-space sampling*

We explored different q-space sampling regimes, scanning at different b-values and here we show results for ex vivo data of the same brain with a range of effective b-values from ~3,000 to ~8,000 s/mm$^2$. Specifically, DW-SSFP data were collected using q-values of 150 and 225 cm$^{-1}$ (effective b-values of 3,200 and 5,700 s/mm$^2$), gradient amplitude of 34.7 mT/m and 52.0 mT/m respectively, gradient duration of 10.16 ms, flip angle of 14 degrees and TR of 21 ms. In addition, data were collected using q-values of 225 and 275 cm$^{-1}$ (effective b-values of 6,200 and 7,900 s/mm$^2$), gradient amplitude of 42.6 mT/m and 52.0 mT/m respectively, gradient duration of 12.42 ms, flip angle of 14 degrees and TR of 26 ms.

Considering these two regimes, we derived quality metrics, including angular CNR and metrics from fibre orientation estimation. We summarise fibre orientation modelling by taking the average of the dispersion for the first fibre orientation and the percentage of voxels with two- and three-way crossings in regions of interest (ROIs). ROIs were hand-drawn in template space and registered to each diffusion space and included the centrum semiovale and midbody of the corpus callosum. We compared how contrast differences translated to increased precision in fibre orientation mapping (reduced uncertainty in the corpus callosum orientations) and increased sensitivity to detecting fibre crossings in the centrum semiovale. We found that increasing the b value from 6,000 s/mm$^2$ and above led to reduced orientation precision and sensitivity in detecting crossings for a spatial resolution of 400 µm (Supplementary Fig. S2). We therefore opted for the lower effective b values of 3,200 and 5,700 s/mm$^2$ for subsequent experiments.

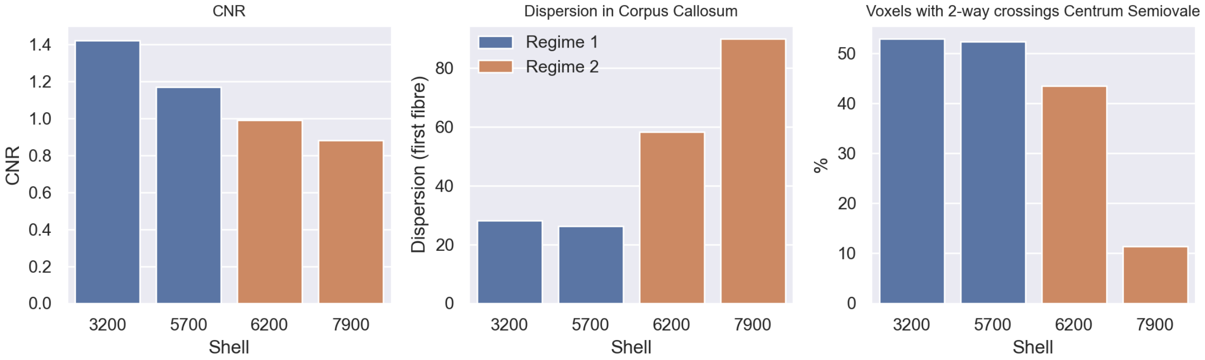


***Supplementary Fig. S2*** *- Assessment of fibre modelling metrics across different DW-SSFP acquisition regimes. Left: contrast to noise (CNR) for each shell. Middle: dispersion in the first fibre population in the mid-body of the corpus callosum. Right: the percent of voxels with 2-way crossings supported by the data in the centrum semiovale.*

#### *Summary of acquisition parameters and sessions*

| | In vivo 750 µm *iso* | In vivo 580 µm *iso* | Ex vivo 400 µm *iso* | Ex vivo 300 µm *iso* |
|---|---|---|---|---|
| Ares | ✓ | ✓ | | |
| Curly | ✓ | | | |
| Gandalf | ✓ | | | |
| Larry | ✓ | | | |
| Moe | | | ✓ | |
| Pear | ✓ | | ✓ | ✓ |

***Supplementary Table S1*** *– Overview of acquisitions to-date.*

| | Spatial res (µm iso) | Matrix | TE (ms) | TR (s) | ES (ms) | b=0 / b=1000 / b=2000 #vols | AP + PA runs | #vols | Scan time |
|---|---|---|---|---|---|---|---|---|---|
| Gandalf* | 750 | 144 × 144 × 80 | 65.6 | 7.35 | 0.333 | 6 / 40 / 39 | 6 + 6 | 1,020 | 2 h 30 min |
| Pear | 750 | 144 × 144 × 68 | 65.8 | 8.26 | 0.333 | 8 / 54 / 53 | 4 + 4 | 920 | 2 h 10 min |
| Curly | 750 | 144 × 144 × 66 | 65.8 | 8.02 | 0.333 | 8 / 54 / 53 | 4 + 4 | 920 | 2 h |
| Larry | 750 | 144 × 144 × 74 | 66.0 | 8.99 | 0.333 | 8 / 54 / 53 | 4 + 4 | 920 | 2 h 20 min |
| Ares (1) | 580 | 282 × 156 × 78 | 79.6 | 9.45 | 0.407 | 8 / 54 / 53 | 4 + 4 | 920 | 2 h 30 min |
| Ares (2)† | 580 | 282 × 184 × 78 | 77.2 | 9.18 | 0.406 | 8 / 54 / 53 | 2 + 2 | 460 | 1 h 10 min |
| Ares (3)† | 750 | 144 × 144 × 74 | 66.0 | 8.99 | 0.333 | 8 / 54 / 53 | 2 + 2 | 460 | 1 h 10 min |

***Supplementary Table S2*** *– Summary of key acquisition parameters for in vivo data. TE = echo time; TR = repetition time; ES = echo spacing. *After Gandalf data were acquired, gradients were replaced. TR was set to slightly longer times to prevent potential gradient over-heating and failure. † Following a scanner repair, Ares data were re-acquired exploring acquisition across resolution using a subset of repeats. The new repeats also have different slice angulation than the first scan.*

| | Spatial res (µm iso) | Effective b-values (s/mm$^2$) | q-values (cm$^{-1}$) | Gradient amplitudes (mT/m) | Gradient duration (ms) | TE (ms) | TR (ms) | Flip angle (degrees) |
|---|---|---|---|---|---|---|---|---|
| Moe | 400 | 0, 3200, 5600 | 32, 150, 225 | 7.4, 34.7, 52.0 | 10.16 | 16 | 21 | 14 |
| Pear | 400 | 0, 2900, 5700 | 50, 175, 275 | 9.6, 33.6, 52.8 | 12.24 | 21 | 26 | 24 |
| Pear | 300 | 0, 3900, 5900 | 67, 200, 266 | 9.6, 38.5, 51.2 | 12.20 | 21 | 27 | 24 |

***Supplementary Table S3*** *– Summary of key acquisition parameters for ex vivo DW-SSFP data. TE = echo time; TR = repetition time.*

*In vivo and ex vivo processing pipeline*

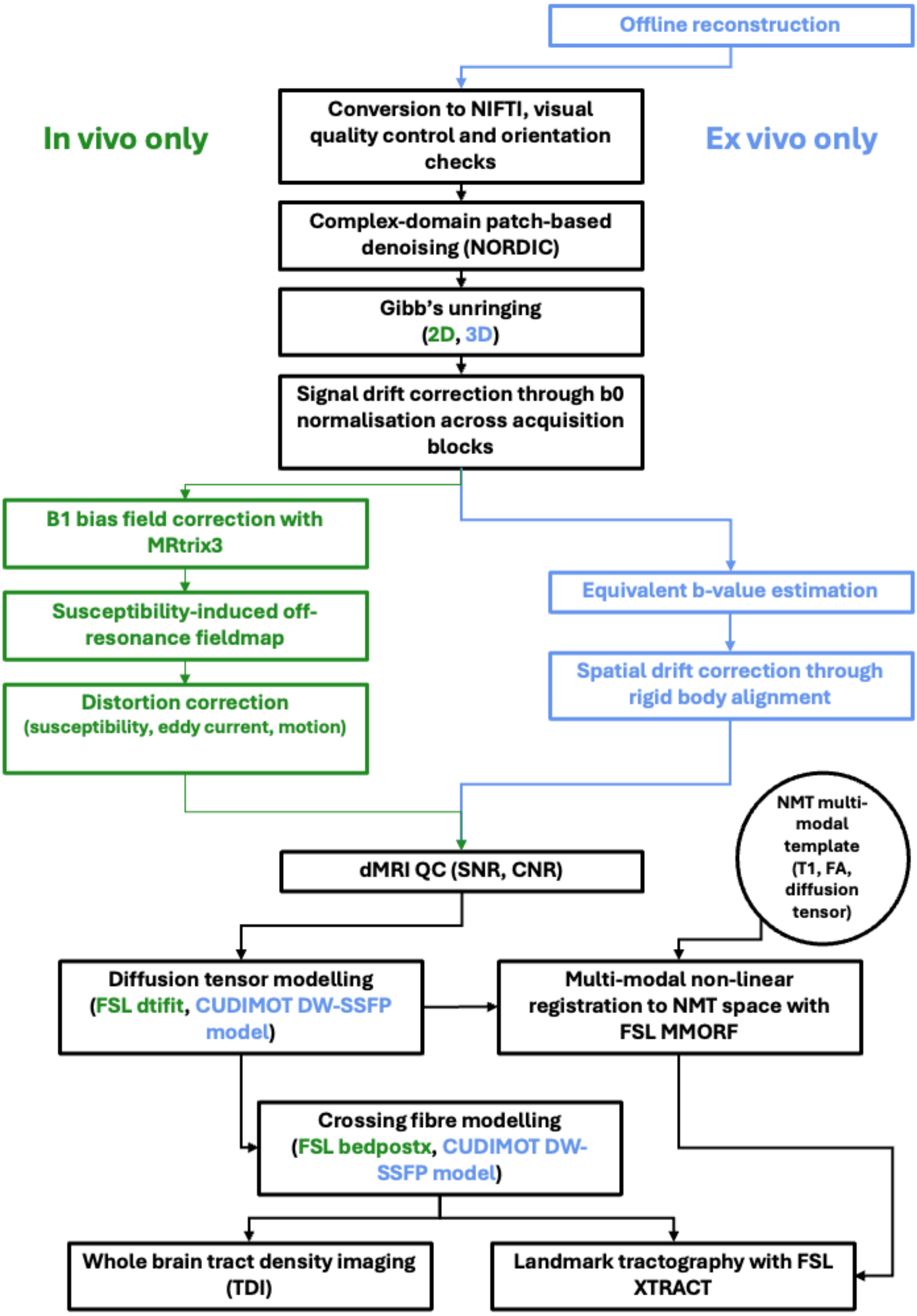


***Supplementary Fig. S3*** *– Schematic overview of the in vivo and ex vivo CMC pipelines. Pipeline steps for only in vivo (ex vivo) data are presented in green (blue). Pipeline steps common for both in vivo and ex vivo data are presented in black.*

#### *Offline reconstruction*

As described, we used custom SENSE1 offline image reconstruction for the ex vivo DW-SSFP data. Using SENSE1 for channel combination minimises the noise floor present in the data. Additionally, some of the scanner reconstructed DW-SSFP data seemed to occasionally suffer from data leakage artefacts (noise bands caused by hardware variable capacitors), which could be dealt with in the offline reconstructed data. Supplementary Fig. S4 examples scanner and custom offline reconstructed data for different volumes from the same acquisition, showing clear gains in signal quality across the brain in the offline reconstruction case.

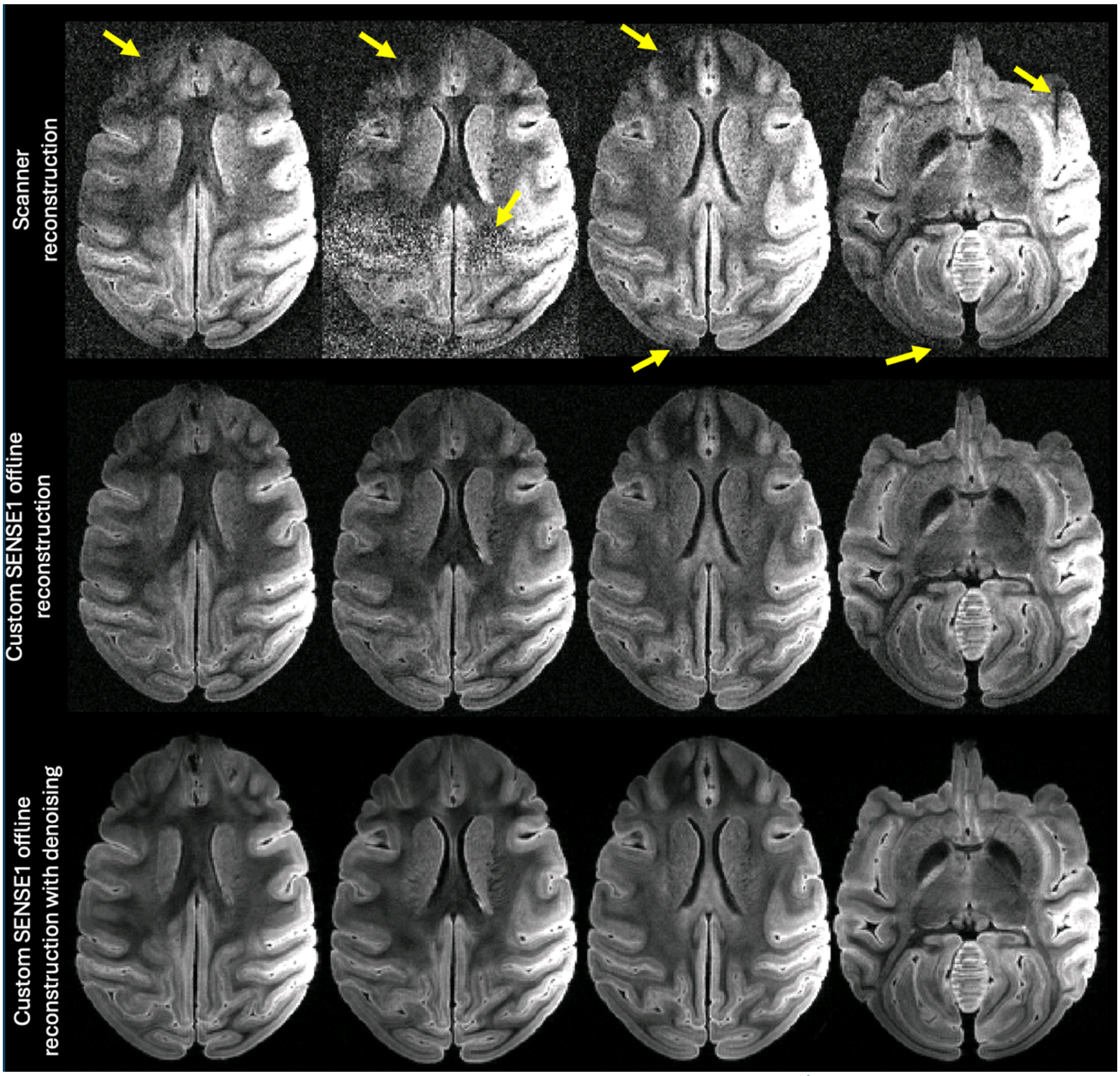


***Supplementary Fig. S4*** *- Examples of ex-vivo DW-SSFP volumes ($b_{eff}$~5,700 s/mm$^2$) reconstructed using the scanner reconstruction engine (top) and the custom offline SENSE1 reconstruction (bottom). Arrows indicate regions of signal dropout and noise bands, which are removed when reconstructed offline.*

#### *Complex-domain image denoising*

Data denoising is performed in the complex domain using NORDIC[3]. We use this approach as previous work has demonstrated reduced noise-induced variance and biases[4]. Supplementary Fig. S5 examples the effect of complex-domain denoising using test ex vivo DW-SSFP data (400 µm resolution) acquired with an effective b-value of approximately 7,800 s/mm$^2$. We observe clear gains in improving signal to noise (SNR) and contrast to noise (CNR) ratios. Further examples are also shown in Supplementary Fig. S4 at a different effective b value.

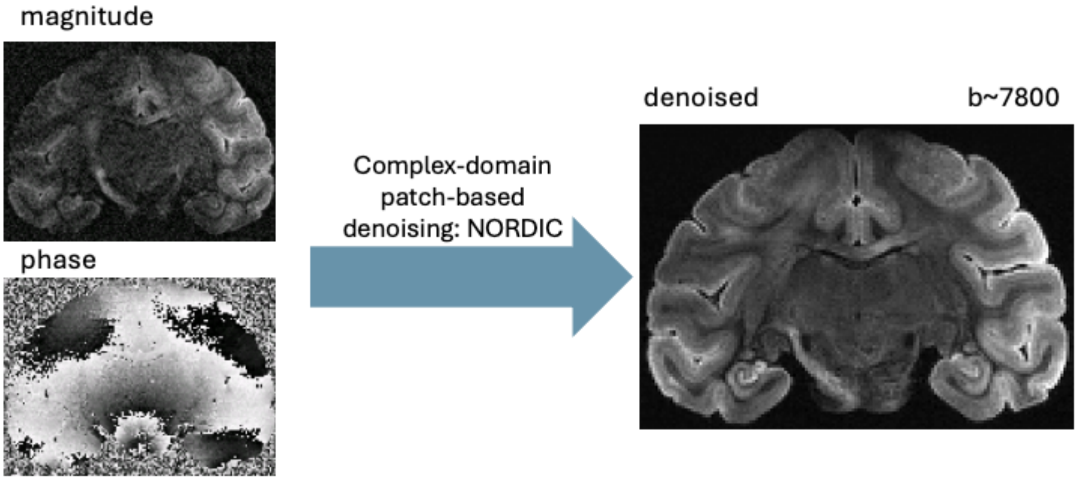


***Supplementary Fig. S5*** *- Complex-domain denoising, exampled using high diffusion-weighted ex vivo data ($b_{eff}$~7,800 s/mm$^2$).*

#### *Gibbs unringing*

For both in vivo and ex vivo, we corrected for Gibbs ringing artefacts (2D for in vivo, 3D for ex vivo) implemented through MRtrix (v3.0.7)[5–7]. We example the effective removal of Gibbs ringing artefacts in Supplementary Fig. S6.

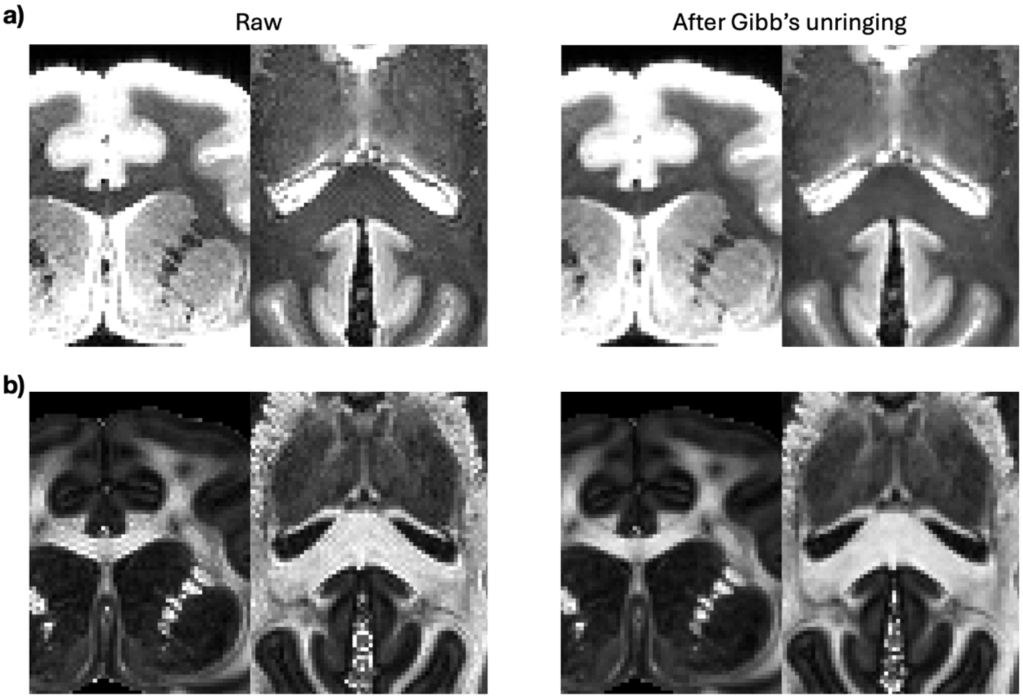


***Supplementary Fig. S6*** *- Example of the removal of Gibbs ringing artefact for the ex vivo data, showing a) a single raw and processed volume zoomed in on the anterior of the brain and b) the downstream effect on fractional anisotropy maps.*

***Signal intensity drift correction***

We assessed mean b=0 signal throughout acquisitions before and after signal drift correction. In both cases, we see that signal drift correction effectively modifies the mean b=0 signal to minimise temporal drift. Signal drift correction is exampled for a single in vivo and ex vivo dataset in Supplementary Fig. S7.

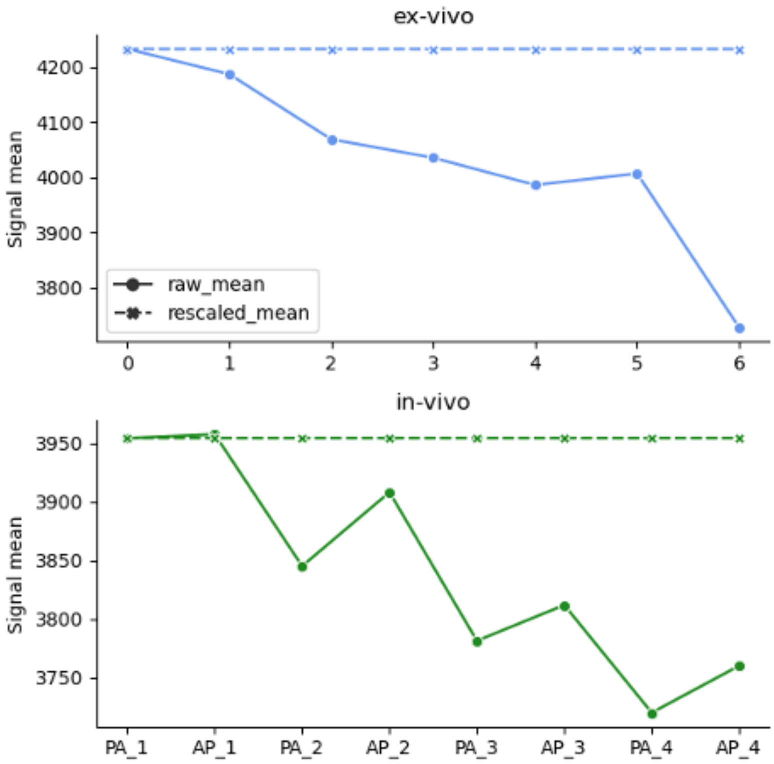


***Supplementary Fig. S7*** *- The mean signal intensities before (solid lines) and after (dashed lines) correction plotted for (top) each b=0 equivalent volume through the ex vivo acquisition and (bottom) averaged across b=0 volumes within each run through the in vivo acquisition.*

#### *Eddy current modelling*

For the in-vivo EPI data, we explored eddy current models, including linear, quadratic and cubic models, evaluated over the four in vivo datasets acquired. We found improvement in alignment between the mean diffusivity maps and anatomical (T2w) data, exampled in Supplementary Fig. S8.

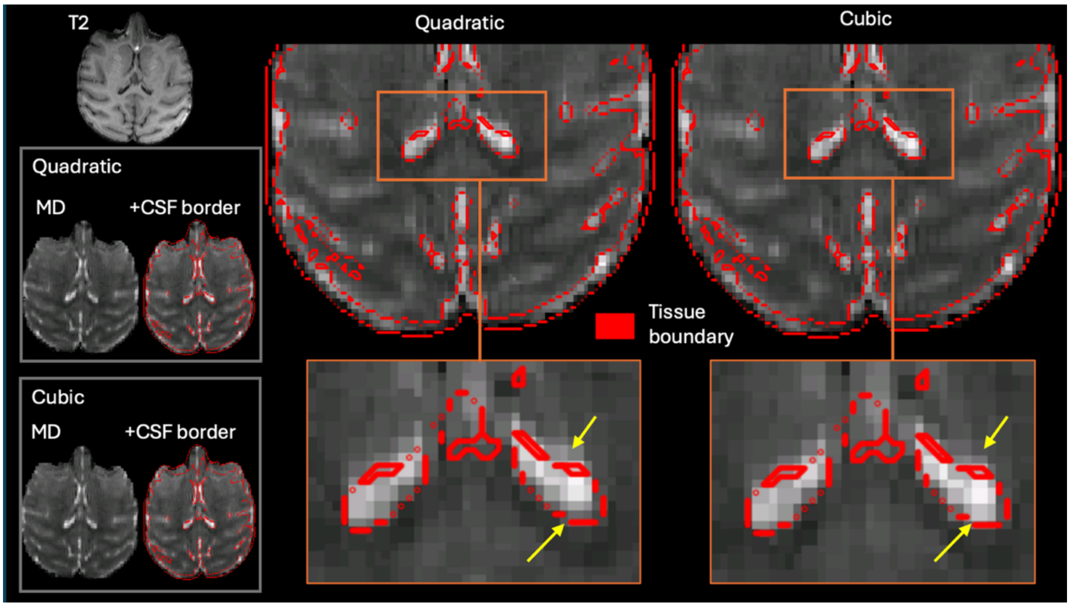


***Supplementary Fig. S8*** *- Qualitative comparisons of quadratic and cubic modelling of eddy currents when correcting eddy current induced distortions. MD maps after quadratic or cubic eddy current correction are overlaid against anatomical boundaries (red outlines).*

We further accounted for large volume-wise displacements, likely present due to temporal drifts over the course of long acquisitions. To account for this, we used smoothing kernels over the iterative estimation of eddy current distortion maps with decreasing full-width half maximum (FWHM: 10, 5, 0, 0, 0 mm over 5 iterations). Supplementary Fig. S9 shows the mean across b=0 volumes, allowing comparison of the default eddy behaviour (five iterations with no smoothing) and the described approach, showing significant improvements in volume alignment, highlighted by the mis-alignment of ventricle signal across volumes (yellow arrow).

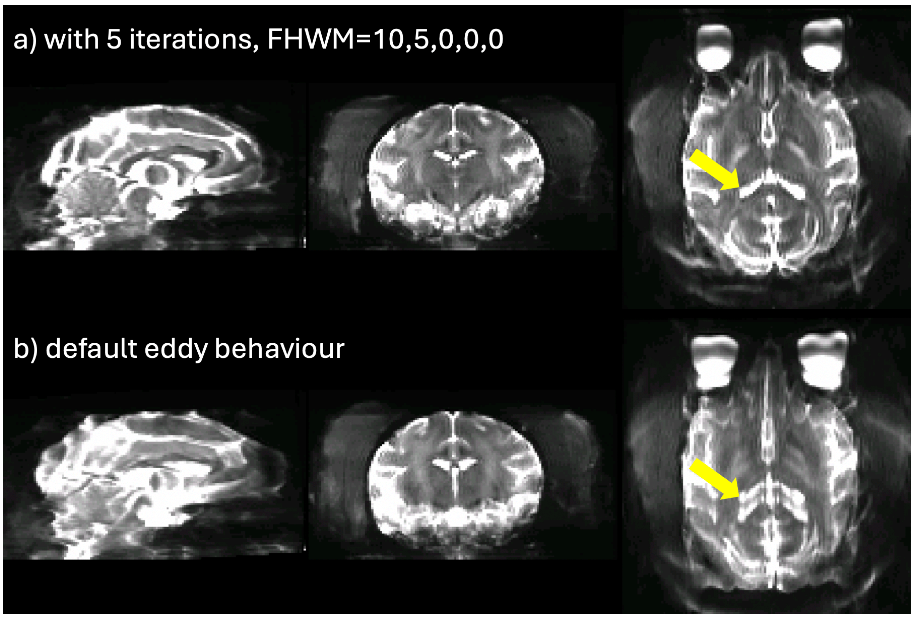


***Supplementary Fig. S9*** *- Comparisons of volume alignment approaches with eddy, using five iterations with varying FWHM (FWHM: 10, 5, 0, 0, 0 mm) smoothing kernels (a) and five iterations with no smoothing (b, default eddy behaviour). The mean of b=0 volumes is calculated for each case and shown. The yellow arrow highlights duplicate ventricles in the default case, arising due to poor alignment of volumes across the acquisition.*

#### *Diffusion tensor maps*

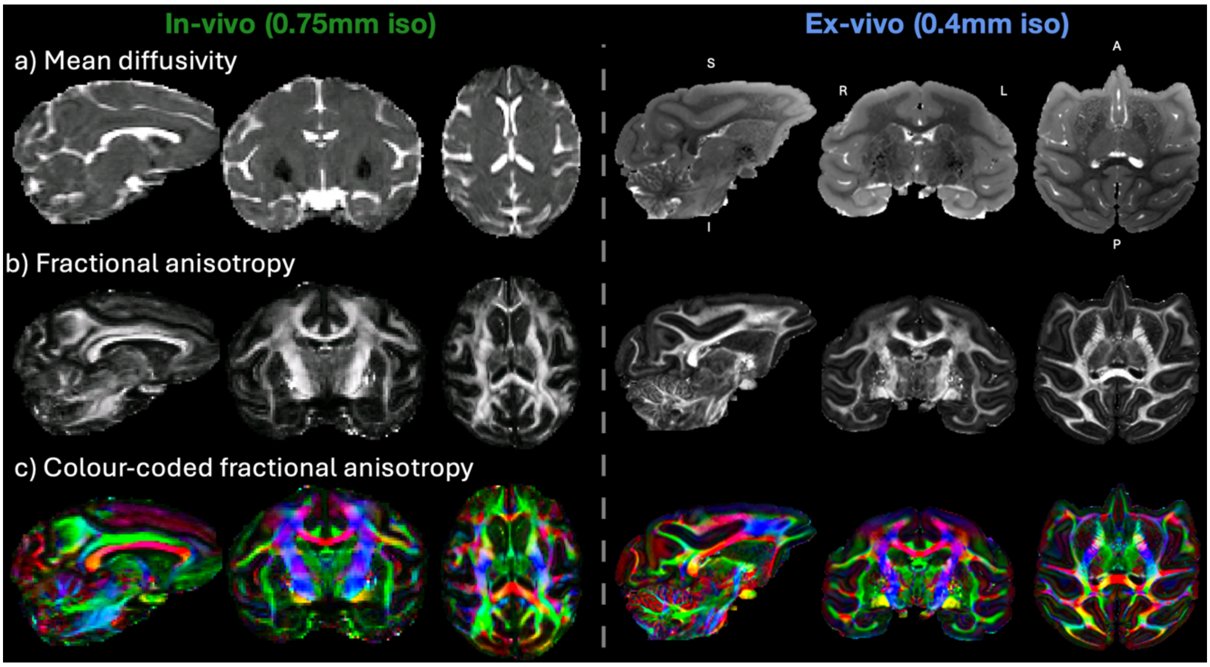


***Supplementary Fig. S10*** *– Example diffusion tensor results for in vivo (left) and ex vivo (right). Top row: mean diffusivity maps. Middle row: fractional anisotropy maps. Bottom row: colour-coded fractional anisotropy maps. The finer spatial resolution of the ex vivo methodology (0.4mm isotropic) can be appreciated compared to that of the in vivo (0.7mm isotropic).*

#### *Multi-modal registration of the same brain to standard space*

Supplementary Fig. S11 shows the NMT-aligned in vivo and ex vivo acquisition for the same animal. Alignment was performed using the non-linear multi-modal registration framework described in the main text.

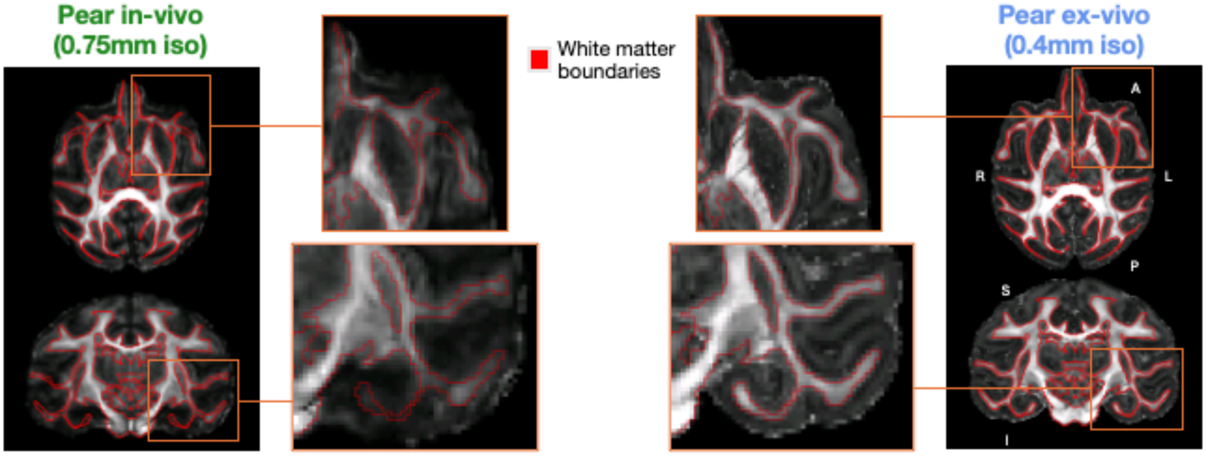


***Supplementary Fig. S11*** *- NMT-space registered in vivo and ex vivo fractional anisotropy (FA) maps from the same brain.*

#### *Fibre Orientation Mapping for DW-SSFP data*

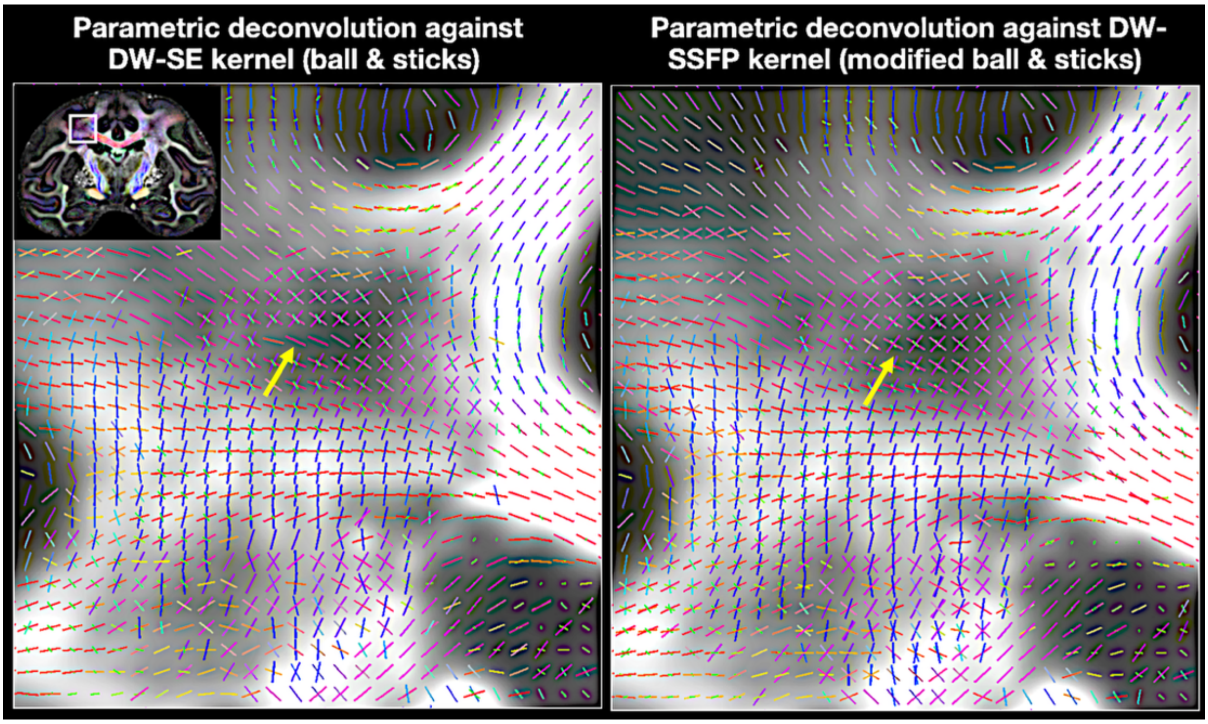


***Supplementary Fig. S12*** *- Fibre orientation estimation for DW-SSFP data using the conventional ball & sticks model (left) with a spin-echo (SE) stick response kernel and the modified ball & sticks model (right) with a DW-SSFP stick response kernel. Higher sensitivity to three-way crossings can be obtained using the latter approach.*

#### *Advancements in imaging resolution in the same animal*

Through hardware, acquisition and processing refinement and optimisation, we were able to push the imaging resolution. We example this below, showing examples of zoomed sections of the same brain imaged across both in vivo (750$\mu m$) and ex vivo (400$\mu m$ and 300$\mu m$) resolutions and another brain imaged at different in vivo resolutions (750$\mu m$ and 580$\mu m$). Clear gains in anatomical detail visible with enhanced resolution are apparent.

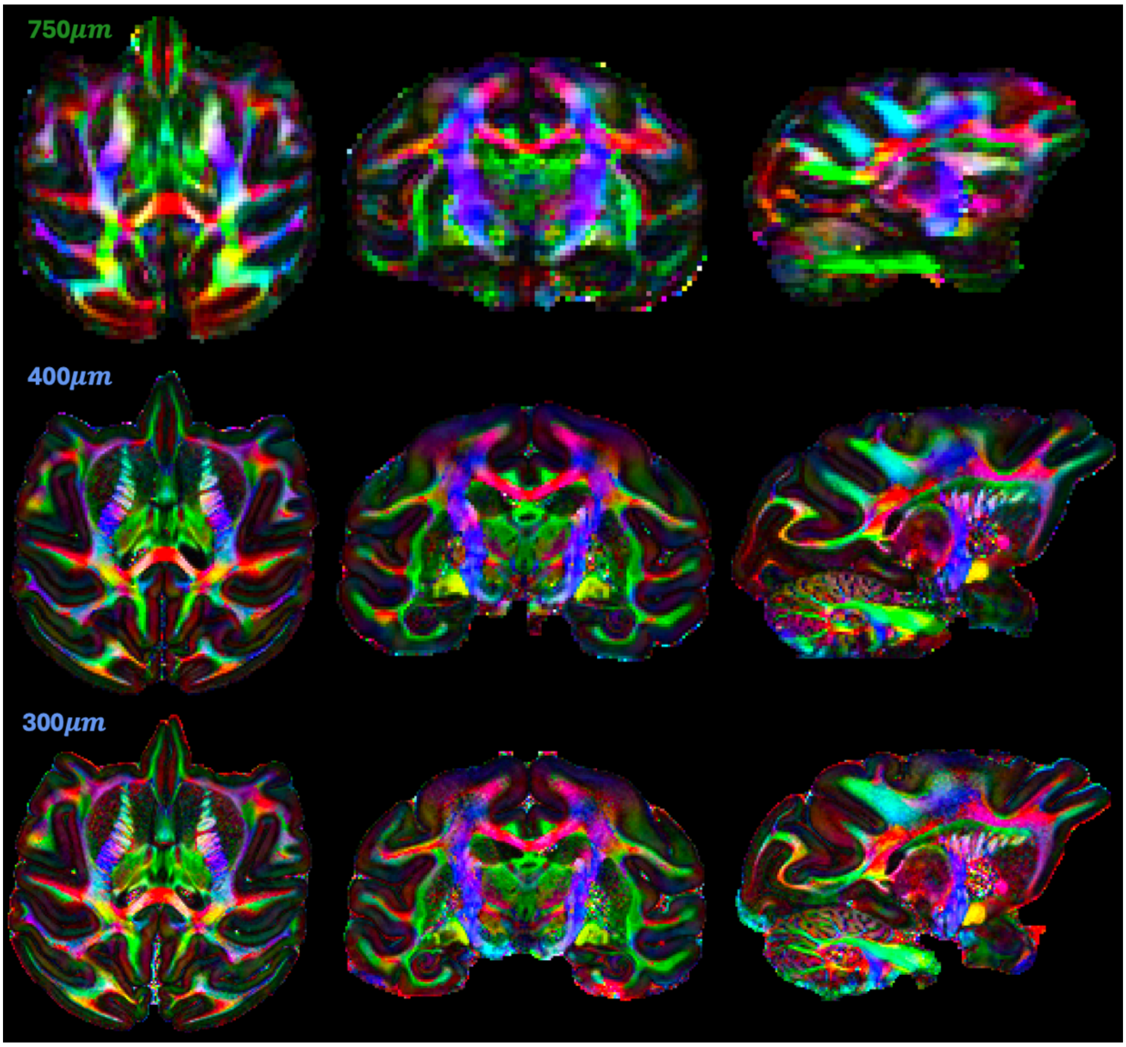


***Supplementary Fig. S13*** *– Examples of the same brain (Pear) imaged across in vivo (750 µm) and ex vivo resolutions (400 µm and 300 µm) showing axial (left), coronal (middle) and sagittal (right) views of the colour-coded fractional anisotropy maps.*

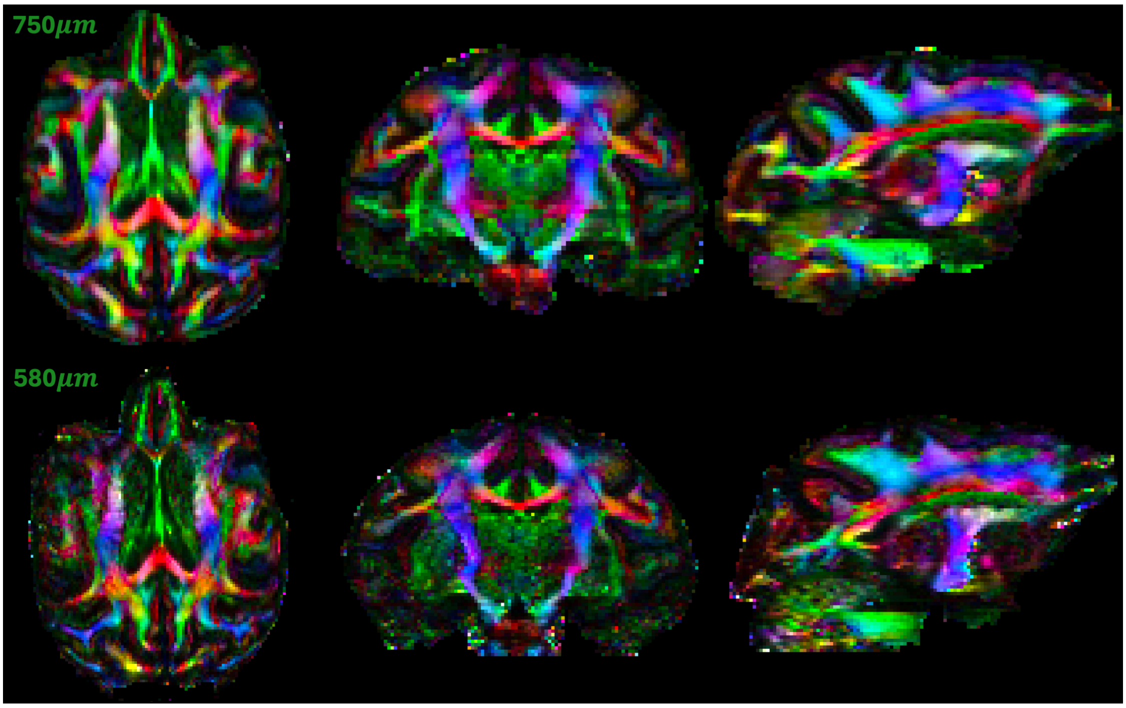


***Supplementary Fig. S14*** *- Examples of the same brain (Ares) imaged across in vivo resolutions (750 µm, 580 µm) showing axial (left), coronal (middle) and sagittal (right) views of the colour-coded fractional anisotropy maps.*

***Track Density Imaging (TDI)***

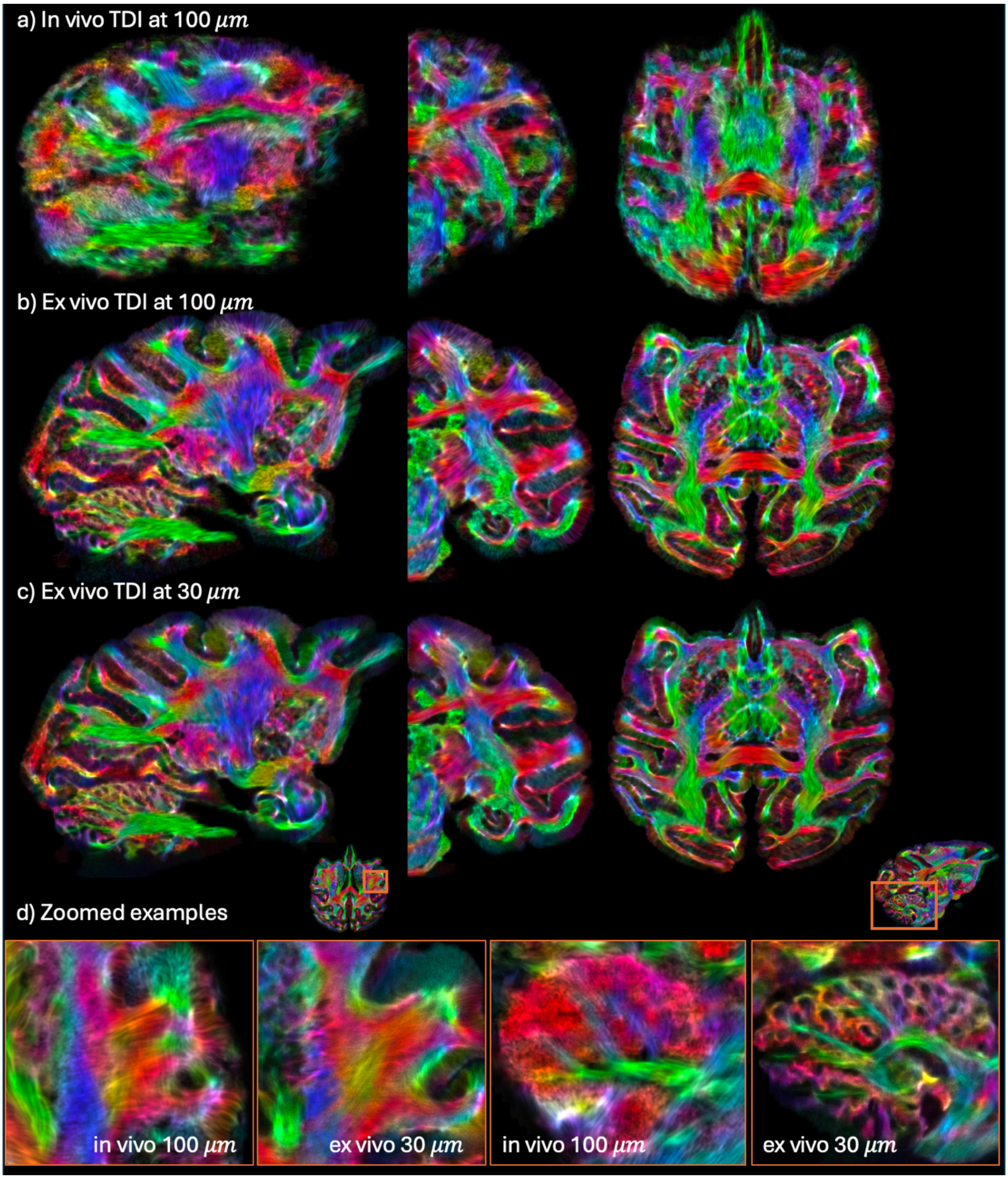


***Supplementary Fig. S15*** *- Track density imaging (TDI) across resolutions, exampled at 100µm in vivo (a) and ex vivo (b), and 30µm (c) ex vivo. Clear benefits are demonstrated when using higher base native resolution, i.e. comparing the same resolution TDI for in vivo (native resolution of 750µm) and ex vivo (native resolution of 400µm) data. Comparing ex vivo TDI resolutions, anatomical detail converges when using TDI resolutions of the order of tens of microns. c) Zoomed in examples of anatomical detail gained comparing 100µm in vivo and 30µm ex vivo TDI.*

***Landmark-based white matter bundle segmentation with XTRACT***

| | Tract | Abbreviation | Bilateral? |
|---|---|---|---|
| **Association Fibres** | Arcuate Fasciculus | AF | Y |
| | Frontal Aslant Tract | FA | Y |
| | Inferior Longitudinal Fasciculus | ILF | Y |
| | Inferior Fronto-Occipital Fasciculus | IFO | Y |
| | Middle Longitudinal Fasciculus | MdLF | Y |
| | Superior Longitudinal Fasciculus I | SLF1 | Y |
| | Superior Longitudinal Fasciculus II | SLF2 | Y |
| | Superior Longitudinal Fasciculus III | SLF3 | Y |
| | Uncinate Fasciculus | UF | Y |
| | Vertical Occipital Fasciculus | VOF | Y |
| **Commissural Fibres** | Anterior Commissure | AC | N |
| | Forceps Major | FMA | N |
| | Forceps Minor | FMI | N |
| | Middle Cerebellar Peduncle | MCP | N |
| **Limbic Fibres** | Cingulum subsection: Dorsal | CBD | Y |
| | Cingulum subsection: Peri-genual | CBP | Y |
| | Cingulum subsection: Temporal | CBT | Y |
| | Fornix | FX | Y |
| **Projection Fibres** | Acoustic Radiation | AR | Y |
| | Anterior Thalamic Radiation | ATR | Y |
| | Corticospinal Tract | CST | Y |
| | Optic Radiation | OR | Y |
| | Superior Thalamic Radiation | STR | Y |
| **Subcortical Fibres** | Amygdalofugal tract | AMF | Y |
| | Extreme capsule (frontal) | EmCf | Y |
| | Extreme capsule (parietal) | EmCp | Y |
| | Extreme capsule (temporal) | EmCt | Y |
| | Muratoff bundle/subcallosal fasciculus | MB | Y |
| | Striatal bundle/external capsule (frontal) | StBf | Y |
| | Striatal bundle/external capsule (parietal) | StBp | Y |
| | Striatal bundle/external capsule (sensorimotor) | StBm | Y |
| | Striatal bundle/external capsule (temporal) | StBt | Y |

***Supplementary Table S4*** *– The full set of 60 white matter fibre bundles reconstructed using XTRACT. Landmark-based tractography protocols have been equivalently defined for both the human and macaque brain, allowing direct comparison across species.*

***Overview of submillimetre macaque diffusion MRI previously published***

| Reference | Spatial resolution (mm) | Number of volumes | b-value (s/mm²) | Acquisition sequence | Field strength (tesla) | Scanner type |
|---|---|---|---|---|---|---|
| (D'Arceuil et al., NeuroImage 2007)[8] | 0.425 | 20 | 4,025 | DW-SE | 4.7 | Preclinical |
| (Wedeen et al., NeuroImage 2008)[9] | 0.512 | 515 | Up to 40,000 | DW-SE EPI | 4.7 | Preclinical |
| (Calabrese et al., Hum. Brain Map. 2014)[10] | 0.4 | 120 | 4,000 | DW-SE | 7 | Preclinical |
| (Thomas et al., PNAS 2014)[11] | 0.25 | 121 | 4,800 | DW-SE EPI | 7 | Preclinical |
| (Calabrese et al., NeuroImage 2015)[13] | 0.15 | 20 | 1,500 | DW-SE | 7 | Preclinical |
| (Azadbakht et al., Cereb. Cortex 2015)[14] | 0.43 | 120 | 8,000 | DW-SE EPI | 4.7 | Preclinical |
| (Donahue et al., J. Neurosci 2016)[15] | 0.43 | 120 | 8,000 | DW-SE EPI | 4.7 | Preclinical |
| (Catani et al., Cortex 2017)[16] | 0.5 | 61 | 4,310 | DW-SE | 4.7 | Preclinical |
| (Folloni et al., 2019)[17] & (Warrington et al., 2020)[18] | 0.6 | 144 | 4,000 | DW-SE | 7 | Preclinical |
| (Saleem et al., NeuroImage 2021)[19] | 0.2 | 112 | Up to 10,000 | DW-SE EPI | 7 | Preclinical |
| (Reveley et al., Nat Commun. 2022)[12] | 0.25 | 126 | 4,800 | DW-SE EPI | 7 | Preclinical |
| (Howard et al., Nat. Commun 2023)[20] | 0.6 | 136 | 4,000 | DW-SEMS | 7 | Preclinical |
| **Current study - CMC** | **0.3** | **112** | **Up to 6,000** | **DW-SSFP** | **10.5** | **Human** |

***Supplementary Table S5** – Overview of **ex vivo** sub-millimetre macaque diffusion MRI studies*

| Reference | Spatial resolution (mm) | Number of volumes | b-value (s/mm²) | Acquisition sequence | Field strength (tesla) | Scanner type |
|---|---|---|---|---|---|---|
| (Janssens et al., NeuroImage 2012)[21] | 0.7 | 256 | 1,000 | DW-SE EPI | 3 | Human |
| (Tounekti et al., NeuroImage 2018)[22] | 0.5 | 30 | 1,000 | DW-SE EPI | 3 | Human |
| (Hayashi et al., NeuroImage 2021)[23] | 0.9 | 500 | Up to 3,000 | DW-SE EPI | 3 | Human |
| (Bihan-Poudec et al., Imaging Neurosci 2023)[24] | 0.4 | 22 | 1,000 | DW-SE EPI | 3 | Human |
| (Valcourt Caron et al., Scientific Data 2025)[25] | 0.7 | 34 | 1,000 | DW-SE EPI | 3 | Human |
| **Current study - CMC** | **0.58** | **464** | **Up to 2,000** | **DW-SE EPI** | **10.5** | **Human** |

***Supplementary Table S6** – Overview of **in vivo** sub-millimetre macaque diffusion MRI studies*

***Supplementary References***


1. Tendler, B. C. *et al.* Diffusion-weighted steady-state free precession imaging in the ex vivo macaque brain on a 10.5T human MRI scanner. 2025.12.12.694017 Preprint at https://doi.org/10.64898/2025.12.12.694017 (2025).
2. Waks, M. *et al.* An RF coil array system for in vivo & ex vivo non-human primate brain studies at 10.5 Tesla. 2025.12.09.692206 Preprint at https://doi.org/10.64898/2025.12.09.692206 (2025).
3. Moeller, S. *et al.* NOise reduction with DIstribution Corrected (NORDIC) PCA in dMRI with complex-valued parameter-free locally low-rank processing. *NeuroImage* **226**, 117539 (2021).
4. Manzano Patron, J. P. *et al.* Denoising diffusion MRI: Considerations and implications for analysis. *Imaging Neuroscience* **2**, 1–29 (2024).
5. Bautista, T., O'Muircheartaigh, J., Hajnal, J. V. & Tournier, J.-D. Removal of Gibbs ringing artefacts for 3D acquisitions using subvoxel shifts. in *Proc. ISMRM* (2019).
6. Kellner, E., Dhital, B., Kiselev, V. G. & Reisert, M. Gibbs-ringing artifact removal based on local subvoxel-shifts. *Magnetic Resonance in Medicine* **76**, 1574–1581 (2016).
7. Tournier, J.-D. *et al.* MRtrix3: A fast, flexible and open software framework for medical image processing and visualisation. *Neuroimage* **202**, 116137 (2019).
8. D'Arceuil, H. E., Westmoreland, S. & de Crespigny, A. J. An approach to high resolution diffusion tensor imaging in fixed primate brain. *NeuroImage* **35**, 553–565 (2007).
9. Wedeen, V. J. *et al.* Diffusion spectrum magnetic resonance imaging (DSI) tractography of crossing fibers. *NeuroImage* **41**, 1267–1277 (2008).
10. Calabrese, E. *et al.* Investigating the tradeoffs between spatial resolution and diffusion sampling for brain mapping with diffusion tractography: Time well spent? *Human Brain Mapping* **35**, 5667–5685 (2014).
11. Thomas, C. *et al.* Anatomical accuracy of brain connections derived from diffusion MRI tractography is inherently limited. *Proc Natl Acad Sci U S A* **111**, 16574–16579 (2014).
12. Reveley, C. *et al.* Diffusion MRI anisotropy in the cerebral cortex is determined by unmyelinated tissue features. *Nat Commun* **13**, 6702 (2022).
13. Calabrese, E. *et al.* A diffusion tensor MRI atlas of the postmortem rhesus macaque brain. *NeuroImage* **117**, 408–416 (2015).
14. Azadbakht, H. *et al.* Validation of High-Resolution Tractography Against In Vivo Tracing in the Macaque Visual Cortex. *Cereb Cortex* **25**, 4299–4309 (2015).
15. Donahue, C. J. *et al.* Using Diffusion Tractography to Predict Cortical Connection Strength and Distance: A Quantitative Comparison with Tracers in the Monkey. *J. Neurosci.* **36**, 6758–6770 (2016).
16. Catani, M. *et al.* Short parietal lobe connections of the human and monkey brain. *Cortex* **97**, 339–357 (2017).
17. Folloni, D. *et al.* Dichotomous organization of amygdala/temporal-prefrontal bundles in both humans and monkeys. *eLife* **8**, (2019).
18. Warrington, S. *et al.* XTRACT - Standardised protocols for automated tractography in the human and macaque brain. *NeuroImage* **217**, 116923 (2020).
19. Saleem, K. S. *et al.* High-resolution mapping and digital atlas of subcortical regions in the macaque monkey based on matched MAP-MRI and histology. *NeuroImage* **245**, 118759 (2021).
20. Howard, A. FD. *et al.* An open resource combining multi-contrast MRI and microscopy in the macaque brain. *Nat Commun* **14**, 4320 (2023).
21. Janssens, T. *et al.* An implanted 8-channel array coil for high-resolution macaque MRI at 3T. *Neuroimage* **62**, 1529–1536 (2012).
22. Tounekti, S. *et al.* High-resolution 3D diffusion tensor MRI of anesthetized rhesus macaque brain at 3T. *NeuroImage* **181**, 149–161 (2018).
23. Hayashi, T. *et al.* The nonhuman primate neuroimaging and neuroanatomy project. *NeuroImage* **229**, 117726 (2021).
24. Bihan-Poudec, Y. *et al.* Cardiovascular effects on high-resolution 3D multi-shot diffusion MRI of the rhesus macaque brain. *Imaging Neuroscience* **1**, imag–1–00039 (2023).
25. Valcourt Caron, A. *et al.* In vivo submillimeter diffusion MRI dataset of 9 macaque brains curated for tractography. *Sci Data* **12**, 1199 (2025).
